# Scale of analysis drives the observed ratio of spatial to non-spatial variance in microbial water quality: insights from two decades of citizen science data

**DOI:** 10.1101/2022.02.01.478743

**Authors:** Daniel L. Weller, Donald E. Weller, Laura K. Strawn, Tanzy M. T. Love

**Affiliations:** Department of Biostatistics and Computational Biology, University of Rochester Medical Center; Department of Environmental and Forest Biology, SUNY College of Environmental Sciences and Forestry; Smithsonian Environmental Research Center, Edgewater MD; Department of Food Science, Virginia Polytechnic University

**Keywords:** Spatial Variation, E. coli, Turbidity, Total Suspended Solids, Bayesian Analysis, Seasonality, Water Monitoring

## Abstract

While fecal indicator bacteria (FIB) testing is used to monitor surface waters for potential health hazards, recent studies report substantial variation in FIB levels and that observed variation appeared dependent on scale of analysis (SOA). Citizen science data and random effects models were used to quantify variance in FIB levels attributable to spatial versus temporal factors. Separately, Bayesian models were used to quantify the ratio of spatial to nonspatial variance in FIB levels, and identify associations between environmental factors and FIB levels. Separate analyses were performed for each stream, lake, and watershed as well as at the state-level (three SOAs). As SOA increased (from waterway to watershed to statewide models), variance attributable to spatial sources generally increased and variance attributable to temporal sources generally decreased. While relationships between FIB levels and environmental factors, such as flow conditions, were constant across SOA, the effect of land cover was both highly dependent on SOA and consistently smaller than the effect of outfalls. This study demonstrates the importance of scale when designing monitoring programs or using FIB monitoring to inform management. Moreover, these data represent a comprehensive survey of water quality in Upstate NY, and the study highlights a valuable publicly available database.

---

Fecal contamination of surface waterways that supply agricultural (e.g., for irrigation, fertigation, dust abatement), drinking, and recreational water has been linked to an increased risk of gastrointestinal and respiratory illness (1–6). For example, multiple *E. coli* O157:H7 outbreaks associated with romaine lettuce between 2017 and 2019 were traced to the probable use of irrigation water contaminated by cattle feces (7–9). Despite this disease burden, pathogen monitoring of agricultural surface water remains impractical due to the number of potential pathogens, the cost of sampling, access to testing laboratories, and methodological limitations amongst other difficulties (10–14). Instead, waterways are monitored for fecal indicator bacteria (FIB) to identify when a presumptive fecal contamination event may have occurred. Regulations and voluntary industry agreements have codified the use of FIB monitoring for assessing surface water suitability for irrigation and recreation as well as for determining if corrective actions are needed [e.g., (15–24)]. For example, Australia and New Zealand, and the United States (US) have established threshold values for coliforms and *E. coli*, respectively, in irrigation water (15, 16), while many countries, including the US, Canada, the European Union, South Africa, and the United Kingdom, have established similar standards for recreational water (17–21).

While FIB-based water standards assume that the proscribed sampling frequency is sufficient to capture temporal variation in water quality, recent studies suggest that the sampling frequencies in several US regulations do not (13, 25–27). As a result, concerns have been raised by scientists and stakeholders about the utility of *E. coli* monitoring programs for assessing microbial hazards in agricultural water (13, 25, 28, 29). Effective implementation of FIB monitoring is further limited by the lack of guidance on the spatial scale at which sampling should be performed. For example, stakeholders have requested guidance on how *E. coli* monitoring outlined in the US Produce Safety Rule should be implemented to adequately capture spatial variation within a water source (29). Additional information on spatial and temporal variation in microbial surface water quality is thus needed to revise current monitoring guidance, especially to determine if changes to monitoring efforts should focus on increased sampling frequency or a larger number of sampling sites. Since recent studies have raised questions about the reliability and efficacy of FIB-based standards for assessing the suitability of waterways for agricultural uses (13, 25, 28), there may be a need for supplementary or alternative indicators. For example, a survey of New York streams proposed using sediment levels (e.g., turbidity) as a supplementary indicator of microbial water quality (13). The repeated identification of a positive association between sediment levels and FIB levels, and between sediment levels and enteric pathogen presence suggests that these associations are reproducible and robust to study design (e.g., location of study, sampling method, laboratory methods (13, 26, 30–33)). However, before incorporating sediment monitoring into existing FIB monitoring programs, an improved understanding of spatial and temporal variation in sediment and FIB levels is also needed (e.g., to inform sampling frequency).

Many surveys have reported considerable variability in FIB and sediment levels over space and time (e.g., (13, 32–38)). For example, Rafi et al. (34) found that *E. coli* levels, and their relationship with watershed size and stream order, varied among three US ecoregions along an East-West gradient (34). Such variability makes sense because surface waterways are hierarchical systems in which water quality is affected by scale-dependent and spatially heterogenous processes and landscapes. Several studies have shown that the strength of associations between FIB levels and land use were dependent on the spatial and temporal scales considered (37, 38). Applied papers have shown that such variability complicates the interpretation of FIB test results and the development of science-based monitoring programs (39). In fact, there are repeated reports that FIB levels differ between samples collected at the same (i) time but from different locations in a waterway (e.g., banks of a river, depths in the water column) or (ii) location but at different times (e.g., within a day, on consecutive days) (13, 40). Given such variability, how should stakeholders use monitoring data to guide decision-making? The real-world implications of these knowledge gaps create a clear need for an improved understanding of temporal versus spatial contributions to variance in microbial water quality. Indeed, while a few papers have discussed how scale of analysis affects observed associations between FIB levels and environmental factors (37, 38), they did not consider the impact of scale on the percent of variance attributable to spatial versus temporal sources, or differences in the impact of scale on observed variability for different water types (e.g., between lakes and streams). These knowledge gaps were highlighted by a review that separately ranked sources of variation in FIB levels in streams, lakes, and coastal systems, but did not make comparisons among water types (39), presumably due to lack of data. The main aims of this study were to address these knowledge gaps by quantifying the impact of spatial scale on the variance in FIB and sediment levels attributable to specific spatial or temporal factors, and on the observed relationship between *E. coli* levels and environmental factors (e.g., land use, weather). A secondary aim was to highlight a publicly-available dataset that can serve as a resource for future studies. Specifically, this study used data from a citizen science initiative run by an NYS certified laboratory that represents one of the more comprehensive publicly-available water quality datasets in Upstate New York State due to the large geographic area sampled (Fig. 1), and the number of water quality parameters measured (Table S1).

**Figure 1:**
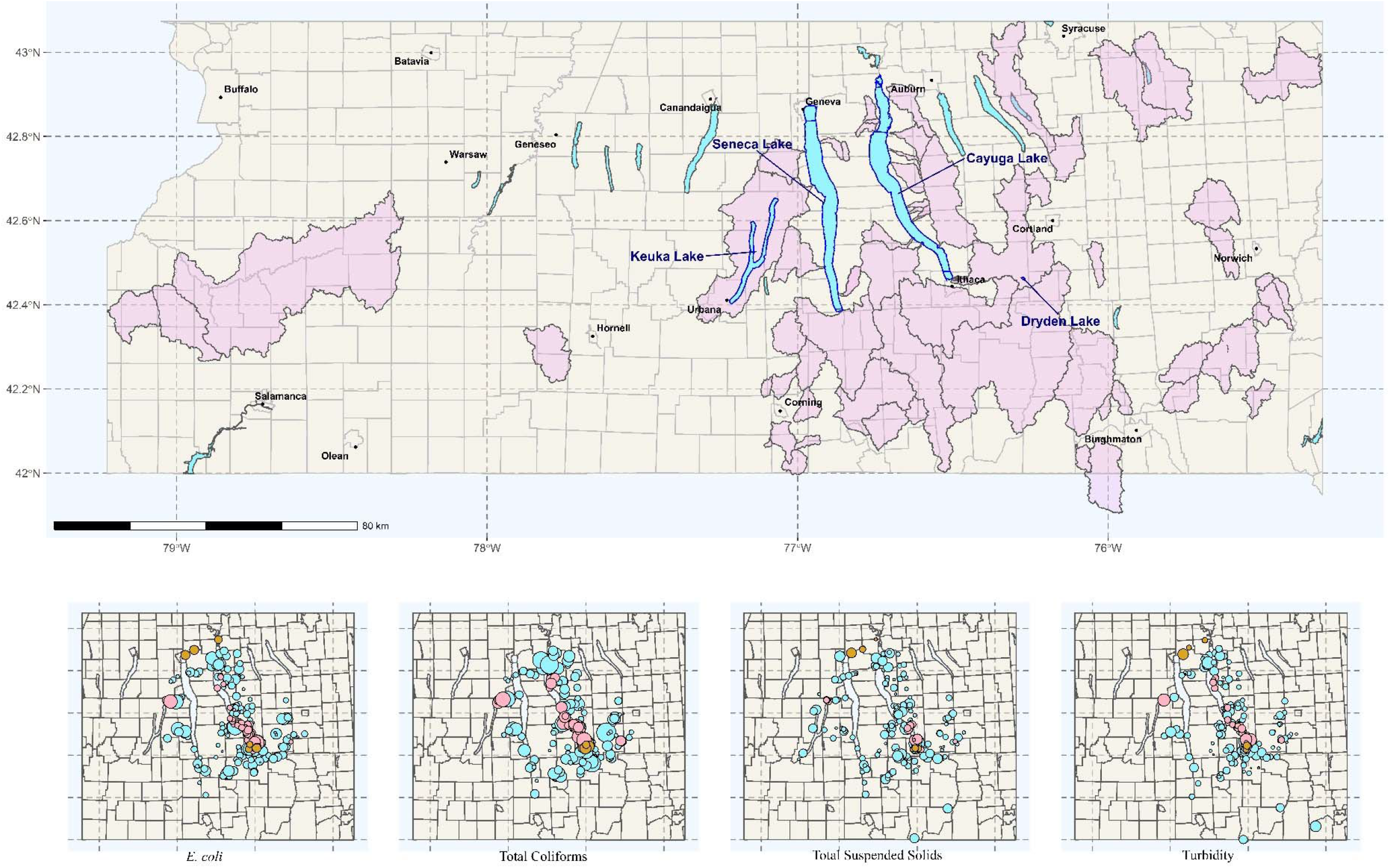
The top map depicts the watersheds for the sampled streams (plum) as well as lakes (blue) and cities (black dots) in the study region. The four maps along the bottom show, from left to right, the average relative concentration of *E. coli*, total coliforms, total suspended solids, and turbidity in samples collected from each canal, lake, and stream site. Larger bubbles indicate higher concentration, smaller bubbles indicate lower concentration. Some samples were not tested for all analytes, so the bottom maps cover a smaller area than the top map. Watersheds omitted from the smaller maps had no data for the four focal parameters of this study.

## Methods

### Study Overview

Data were collected by the Community Science Institute (CSI) and are available at their website http://www.communityscience.org/. While this is a publically-available dataset, this is the first publication to summarize all water quality parameters analyzed (Table S1) and to assess spatiotemporal variation in key parameters. CSI runs a NY Department of Health (NYDOH) certified lab, and the methods used to enumerate each of the parameters reported here are detailed on their website, in their certifications (available from the NYDOH), and in Table S1. Data were collected between 2002 and 2020 (N=11,351 samples) and represent 58 waterways (Fig 1).

### Metadata Collection

For each sample collected, CSI characterized flow conditions as baseflow or stormflow. The presence of outfalls at or immediately upstream of the sampling site, if the site was at the stream mouth (i.e., where the stream entered a lake), and water type (e.g., canals, lake, stream) were also noted. For streams, the inverse-distance weighted proportion of each stream’s watershed under cropland, developed, forest-wetland, and pasture cover was assessed as previously described (41).

Upstream land use was not characterized for canals because they lack consistent, unidirectional flow, and delineating upstream watersheds was not possible. Nor was land use characterized for the sampled lakes (e.g., Cayuga Lake, Seneca Lake), which were sufficiently large that we did not expect land cover in the watershed to be associated with water quality at individual sampling points, particularly when samples were collected away from shore. The county and HUC-4 watershed where each sample was collected were also recorded. However, due to the number of samples collected in the Finger Lakes HUC-4 watershed, samples collected there were further subdivided into the areas draining into Cayuga Lake, Keuka Lake, and Seneca Lake, respectively. We call these subdivisions “watersheds”, and refer to individual streams and lakes as “waterways” to avoid confusion. Additionally, the season, day of the week, and week of the year (i.e., number of weeks since Jan 1^st^) for each sampling event was recorded.

### Descriptive Analysis

Watershed delineation was performed in ArcMap (42), while all other analyses were performed using R version 3.5.3 (43). While 11,351 samples were collected, not all analytes were enumerated in all samples resulting in substantial missingness (Table S1). Four water quality parameters [*E. coli*, total coliforms, total suspended solids (TSS), and turbidity] were selected as the foci of analyses. These parameters were selected because they are relevant to environmental, human, and/or economic health, and they were measured in more than 1,000 samples collected from multiple watersheds. In comparison, fecal coliform levels, another FIB used in monitoring programs, were only enumerated in 155 samples representing three streams, and as such was not included among the focal parameters in the present study. Spatial and temporal patterns for each of the four focal parameters were visualized, as were differences in water quality between baseflow and stormflow conditions, watersheds, and water types. Spearman’s correlation between all continuous parameters (e.g., water temperature, week of the year, year, *E. coli* levels) was assessed and visualized using the corrplot package (44). For the correlation analysis and all other analyses described below the four parameters were log_10_-transformed.

### Statistical Analysis

To assess the effect of scale on observed sources of variance, random effects models and intercept-only Bayesian models of coregionalization (MCOR) were implemented in the following five ways:

- separately for each stream and lake (the “waterway models”)
- separately for each watershed (the “watershed models”)
- using all stream data combined (the statewide “all-stream model”)
- using all lake data combined (the statewide “all-lake model”),
- using all the data (the “statewide model”).

As a result, water type-specific and water type-agnostic comparisons could be made across three scales of analysis (waterway, watershed, and statewide). Multivariable MCOR were implemented in the same five ways to assess the effect of scale on the observed relationship between FIB levels and environmental factors and allow similar comparisons to be made between spatial scales and water types. Five streams (referred to as the direct streams), Deans Creek, Lake Ridge Creek, Mill Creek, Paines Creek, and Townline Creek, were treated as the same waterway for sampling purposes by CSI; as a result, separate waterway models were implemented for each of the five streams as well as all direct streams combined.

In addition to streams and lakes, CSI collected samples from canals/flood control channels (henceforth called canals), springs, ponds, and wetlands. Wetland samples were not used in any analyses here. Spring data was only used in the statewide multivariable MCOR to allow comparison between water types. All sampled canals linked two lakes, drew water from a lake, served as a flood control channel for a lake, or connected a lake to another waterway. Therefore, canal data were grouped with the corresponding lake for the individual lake and all-lake models but kept separate in the descriptive analyses to facilitate visualization of differences between canals and lakes. Results from individual pond and lake models are visualized together in the graphical summaries due to the small number of ponds (N=2). Similarly, pond data were pooled with lake data for the descriptive analysis and all-lake models. As a result, dummy variables for water type were included as covariates to quantify differences among canals, lakes, ponds, streams, and/or springs in the statewide multivariable MCOR.

#### Random Effects Models

Separate random effects models were built to assess the percent of variance in each water quality parameter that could be attributed to specific spatial and temporal factors. Since the aim of the random effects models was to compare variance attributable to spatial versus temporal sources, individual waterway models were not implemented for waterways where all samples were collected from a single site: Chapel Pond, Lake Ridge Creek, Glen Eldridge Creek, or Townline Creek. Data for Lake Ridge and Townline Creek were used in the Direct Streams waterway model, while data for all four waterways were used in the appropriate watershed and statewide models. Each random effects model included one of the following factors as a random effect: month, season, week day (e.g., Monday, Tuesday), week of the year (i.e., weeks since Jan. 1^st^), year, sampling site, and county. For waterways and watersheds where different water types were sampled (e.g., impoundments within streams) or where stream mouths were sampled, random effects models were implemented with a random effect of water type. Models were fit using the lme4 package (45), and *R*^2^ was calculated using the MuMin package (46) as previously described (47). Results for each water type (stream versus lake) and each scale of analysis were visualized using ggplot (48).

#### Bayesian Models of Coregionalization (MCOR)

Intercept-only MCOR were fit to characterize the ratio of spatial to non-spatial variance for each water quality parameter (*E. coli*, total coliforms, TSS, and turbidity), and to quantify the effective range, which is the distance at which FIB and sediment levels became independent. Since the aim of the intercept-only models was to quantify the ratio of spatial to non-spatial variance, individual waterway models were not implemented for the four waterways where all samples were collected from a single site as discussed above. Separately, multivariable MCOR were fit to identify drivers of elevated or reduced *E. coli* levels, while explicitly accounting for spatial autocorrelation among sites. Multivariable models were only developed for *E. coli* because *E. coli* monitoring is commonly used to identify when a presumptive fecal contamination event has occurred in agricultural and recreational waterways, and multiple governments have established *E. coli*-based standards for agricultural and recreational waterways (11, 16, 17, 29, 49). As such, understanding drivers of elevated *E. coli* levels is of economic, environmental, and human health importance (16, 17, 29, 49).

Before developing the univariable and the multivariable MCOR, semivariograms were fit to help generate informative priors for phi (3/effective range), tau (nugget, or non-spatial variance), and rho (partial sill, or spatial variance). If the semivariogram could not be fit using an exponential covariance model then a Gaussian model was attempted; if a semivariogram could still not be fit then non-informative priors were set (phi=3/330, nugget=semivariance, partial sill=semivariance/5). The MCOR were fit and estimates were quantified using the spBayes and coda packages (50–52). Ten thousand iterations were performed for each MCOR, but the first 7,000 were discarded as burn-in. For all MCOR, the 25^th^, 50^th^, and 75^th^ percentiles for phi, tau, rho, the ratio of spatial to non-spatial variance, and the percent of variance attributable to spatial sources were calculated. For the multivariable MCOR the 25^th^, 50^th^, and 75^th^ percentiles for the effect estimates were also quantified.

In all multivariable MCOR, the following factors were included as explanatory factors when applicable: log_10_ TSS; flow conditions (baseflow=reference-level); month (April= reference-level); if the sampling site was at the stream mouth (No= reference-level); if the site was at or immediately downstream of an outfall (No= reference-level); and year. In models that included stream but not lake data, between one and three land cover parameters (IDW proportion of area under crop, developed, or pasture cover) were also included. The number of land cover factors depended on the correlations among the three cover parameters for the subset of data used in the model. Developed cover was only included if the correlation between developed and pasture cover was <0.7, while cropland was only included if the correlation of developed with crop and pasture cover, and of pasture with crop cover were <0.7. Forest-wetland cover was not included in any models because the correlation between forest-wetland cover and one or more of the other land covers was >0.7 for all data subsets. In the statewide model and all-lake models, water type was also included as a covariate to see if *E. coli* levels varied substantially among water types. Similarly, watershed was included in the statewide model to see if *E. coli* levels varied substantially among watersheds.

## Results

Between 2002 and April 2020, the Community Science Institute (CSI) and its volunteers collected and tested 11,351 surface water samples (Table S1). The waterbodies sampled spanned Upstate New York from Lake Erie to the Capital Region, and Lake Ontario to the Pennsylvania border (Fig 1). The number of samples increased from 18 samples collected in 2002 to a peak of 1,310 samples in 2014 (Fig S1). Following sample collection, microbe (e.g., *E. coli*, total coliforms), nutrient (e.g., nitrate, total phosphorous), heavy metal (e.g., arsenic), and/or sediment (e.g., TSS, turbidity) concentrations as well standard physicochemical parameters (e.g., conductivity, pH) were enumerated. However, each parameter was only measured in a subset of the 11,351 samples collected (Table S1). This paper focused on four parameters, *E. coli*, total coliforms, total suspended solids (TSS), and turbidity (Table 1; Table S1; Fig 1).

**Table 1:** Number of samples tested for the four focal water quality parameters of the present study collected from each water type and overall.

|  | Lakes <sup>a</sup> |  | Streams |  | Total |  | Enumeration Method |
| --- | --- | --- | --- | --- | --- | --- | --- |
|  | Samples | Sites | Samples | Sites | Samples | Sites |  |
| *E. coli* | 1,738 | 92 | 4,025 | 158 | 5,763 | 250 | EPA Method 1604 <sup>b</sup> |
| Total Coliforms | 1,166 | 78 | 3,375 | 148 | 4,541 | 226 | EPA Method 1604 <sup>b</sup> |
| Total Suspended Solids | 935 | 50 | 3,864 | 153 | 4,799 | 203 | SM 2450 D-2011 <sup>c</sup> |
| Turbidity | 1,128 | 57 | 3,742 | 156 | 4,870 | 213 | EPA Method 180.1 <sup>d</sup> |

<sup>a</sup> In addition to streams and lakes, the CSI collected samples from canals/flood control channels and ponds. All sampled canals linked two lakes, drew water from a lake, served as a flood control channel for a lake, or connected a lake to another waterway. Therefore, canal data were grouped with the lake data. Due to the small number of ponds (N=2), pond data were pooled with lake data as well. In the present table, 114, 99, 113, and 115 of the “lake” samples correspond to samples collected from canals/flood control samples and used for *E. coli*, total coliform, total suspended solids, and turbidity enumeration, respectively. Eight of the “lake” samples correspond to samples collected from a pond and used for *E. coli*, total coliform, total suspended solids, and turbidity enumeration.
<sup>b</sup> https://www.epa.gov/sites/default/files/2015-08/documents/method_1604_2002.pdf
<sup>c</sup> https://www.standardmethods.org/doi/10.2105/SMWW.2882.030
<sup>d</sup> https://www.epa.gov/sites/default/files/2015-08/documents/method_180-1_1993.pdf

### Visualizing Spatial and Temporal Patterns

Temporal patterns were assessed visually using loess-smoothed trendlines at three temporal scales: interannual, intra-annual (or seasonal), and event-scale (i.e., stormflow versus baseflow). Because sampling was not conducted at the same site or on the same waterway on consecutive days or weeks, day-to-day and week-to-week variation were not assessed. Based on the loess-smoothed trendlines, there was no evidence of substantial interannual variation in TSS and turbidity levels (Fig S3). Conversely, *E. coli* levels appeared to increase, on average, between 2015 and 2020 in streams and lakes but not canals, while total coliform levels appeared to decrease in streams and canals over the same period (Fig S3). Microbial levels in lakes followed a sinusoidal pattern with levels generally increasing from 2015 to 2020 (Fig S3). *E. coli* and total coliform levels also showed evidence of seasonality (Figs S2–S3) with levels peaking between May and October; this trend was more pronounced in canal and lake samples compared to stream samples. Conversely, TSS and turbidity peaked in the colder months (Fig S2–S3). Both microbial and sediment levels were substantially higher under stormflow than baseflow conditions (Fig 2).

**Figure 2:**
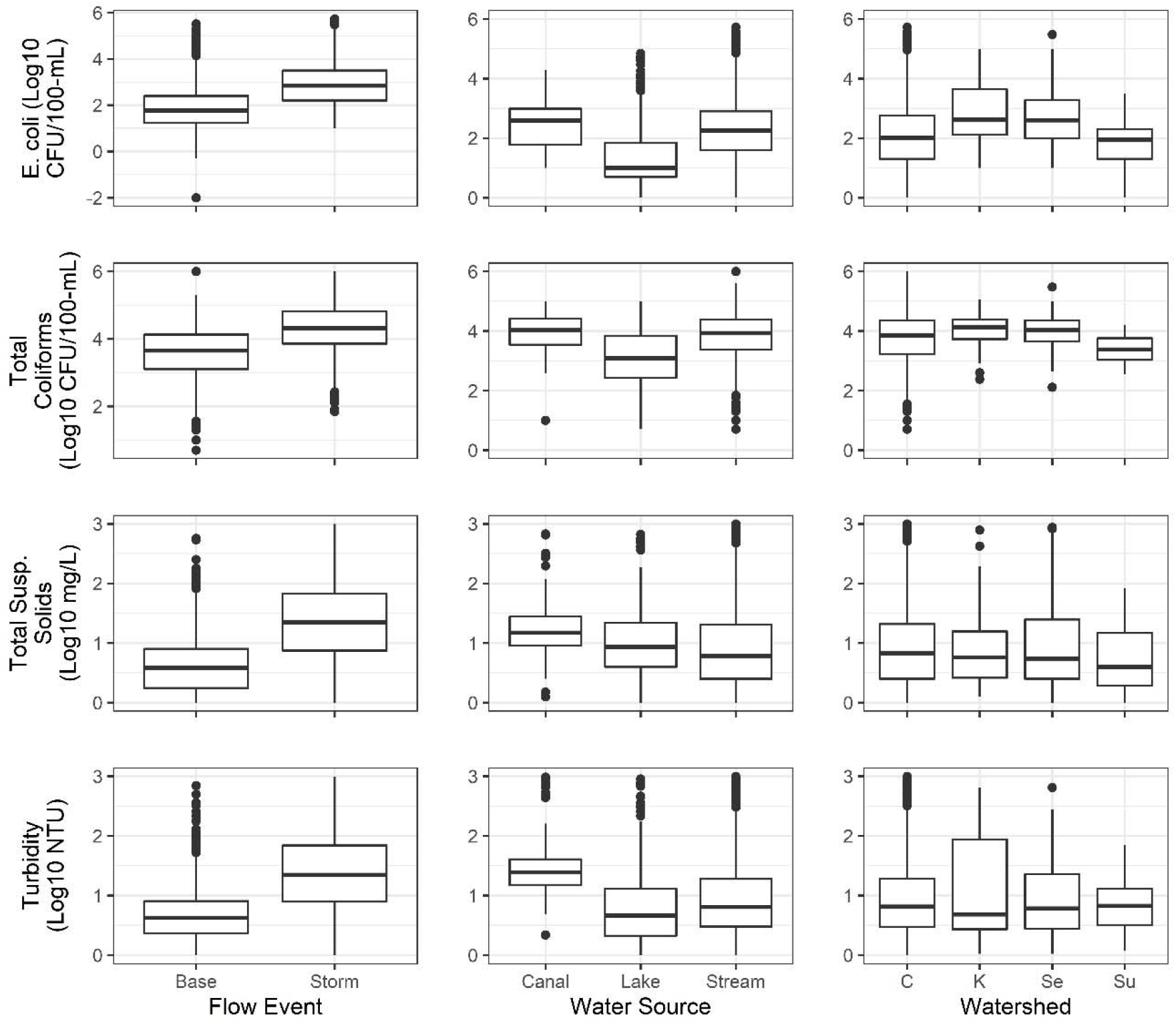
Distribution of log10 *E. coli*, total coliform, total suspended solids, and turbidity for baseflow versus stormflow events, each type of water source, and each HUC-4/Finger Lake watershed providing data for one or more of the four focal water quality parameters (C = Cayuga Lake, K = Keuka Lake, Se = Seneca Lake, Su = Susquehanna River).

Spatial patterns were examined visually by comparing the distribution of each parameter (*E. coli*, total coliforms, TSS, and turbidity) between water sources and watersheds (Fig 2), and by plotting parameter concentration against latitude, longitude, and land cover (Fig S4). On average, all parameters decreased from North-to-South possibly because of North-South land-use patterns since trends in land cover mirrored trends in water quality (Fig S4). Specifically, southern regions were heavily forested while northern regions were largely agricultural (Fig S4). While sediment levels did not appear to differ substantially between watersheds *E. coli* levels were substantially higher in the Keuka and Seneca Lake watersheds compared to the Cayuga Lake and Susquehanna River watersheds (Fig 2). Both *E. coli* and total coliforms levels were substantially higher in canals and streams compared to lakes (Fig 2). Sediment levels were similar in lakes and streams but higher in canals; in fact, the interquartile range (IQR) for turbidity levels in canals did not overlap the IQR for turbidity levels in either lakes or streams (Fig 2).

### The percent of variance attributable to spatial versus nonspatial sources was dependent on the scale of analysis, water type, and waterway/watershed

Random effects models were used to characterize the percent of variance in each water quality parameter accounted for by different spatial (e.g., site, waterway, watershed) and temporal (e.g., week day, week of the year, year) factors as well as water type (Fig. 4, Table S2–S3). Separately, intercept-only MCOR models were used to quantify the percent of variance attributable to spatial sources for each parameter (Fig. S5, Table S2). Comparisons between scales of analysis in the variance attributable to a given source were visualized using boxplots. Briefly, when the point representing the variance attributable to the statewide, all-lake or all-stream models fell outside the interquartile range (IQR) for the waterway or watershed models or when the IQRs did not overlap for the waterway and watershed models it indicated a substantial difference in the variance accounted for by the given factor between model types. Similarly, there was evidence of a potential difference if the median of one group was outside the IQR for the another. Lastly, the difference between the first and third quartile (bottom and top of the box) provides insight into the data distribution, with shorter boxes meaning data hovers around the median values and larger boxes indicating greater variability.

In general, there was considerable variation in the percent of variance attributable to spatial versus non-spatial sources and in the variance attributable to any given factor among the waterway and watershed models (Fig. 4, S5; Table S2, S3) suggesting differences in sources of variance between waterways and between watersheds. However, the variance in all four water quality parameters attributable to spatial factors, such as site, was generally higher for the statewide and watershed models than the waterway models (Fig 4, Table S2–S3). The variance attributable to site in the statewide, and the all-lake and/or all-stream models fell outside the IQR for the waterway models for all four parameters, and, for turbidity and TSS, the IQR for the watershed models did not overlap the IQR for the waterway models (Fig 4). Conversely, the variance attributable to temporal factors largely decreased as the spatial scale of analysis increased (Fig. 4, Fig S5, Table S3). A similar pattern was evident in the intercept-only MCOR models for *E. coli* and total coliforms but not TSS or turbidity. For total coliforms, the IQR for the percent of variance attributable to spatial factors in the watershed MCOR models was higher and did not overlap the IQRs for the lake and/or stream models (Fig S5; Table S2). While a similar pattern was observed for *E. coli*, the IQR, but not the median, for the watershed models overlapped the IQR for the stream models (Fig S5).

While the variance in all four parameters attributable to spatial factors increased as the scale of analysis increased, variance attributable to temporal factors decreased as scale increased and was generally lower in models built using the lake as opposed to stream data. The IQR for the variance =attributable to month, week day, and week of the year in the waterway and watershed models were generally above the variance attributable to the given factor in the statewide, all-lake, and all-stream models (Fig 4; Table S5). For example, the IQR for the variance attributable to week of the year in the stream models did not overlap the IQR for the watershed or lake models for *E. coli*, TSS, and turbidity (Fig. 4; Table S5). Similar differences across scales and/or between water types were observed for *E. coli* in the variance attributable to year, for total coliforms in the variance attributable to month and year, for TSS in the variance attributable to month, week day, and year, and for turbidity in the variance attributable to week day.

Across all four parameters, less variance was attributable to year than to temporal factors that account for seasonality (month, season, and week of the year; Fig. 4, Table S3). Indeed, the temporal factors that accounted for the greatest variance in microbial water quality were seasonal (month, season, or week of the year; Fig 3; Table S3) for all models except one watershed and two stream *E. coli* models for as well as one stream and two lake coliforms models. Overall, variance attributable to season appeared to decrease as scale of analysis increased and was lower for lake compared to stream models (Fig 3; Table S3). Although substantial variance in microbial water quality was attributable to season, little variance was attributable to year for all scales of analysis and water types. While year accounted for 0% of variance in one watershed, three stream, and two lake *E. coli* models, the smallest variance accounted for by week of the year was 8% (Table S3). When the ratio of variance attributable to week of year to variance attributable to year was calculated for each model (replacing 0% with 1%), the ratio was between 1.4 and 25.0 (median=5.9) for watershed models, 1.1 and 77.0 (median=4.5) for stream models, and 2.0 and 23.0 (median=10.9) for lake models. Additionally, 3.7, 3.7, and 3.4 times as much variance was attributable to week of the year compared to year in the statewide, all-stream, and all-lake models. It is also interesting to note that the amount of variance attributable to spatial factors, like site, compared to temporal factors, such as week of the year, also differed by water type. For instance, site and week of the year accounted for 50% and 17%, respectively, of variance in *E. coli* levels in the all-lake model, but for 23% and 22%, respectively, of variance in *E. coli* levels in the all-stream model (Fig. 4, Table S3).

**Figure 3:**
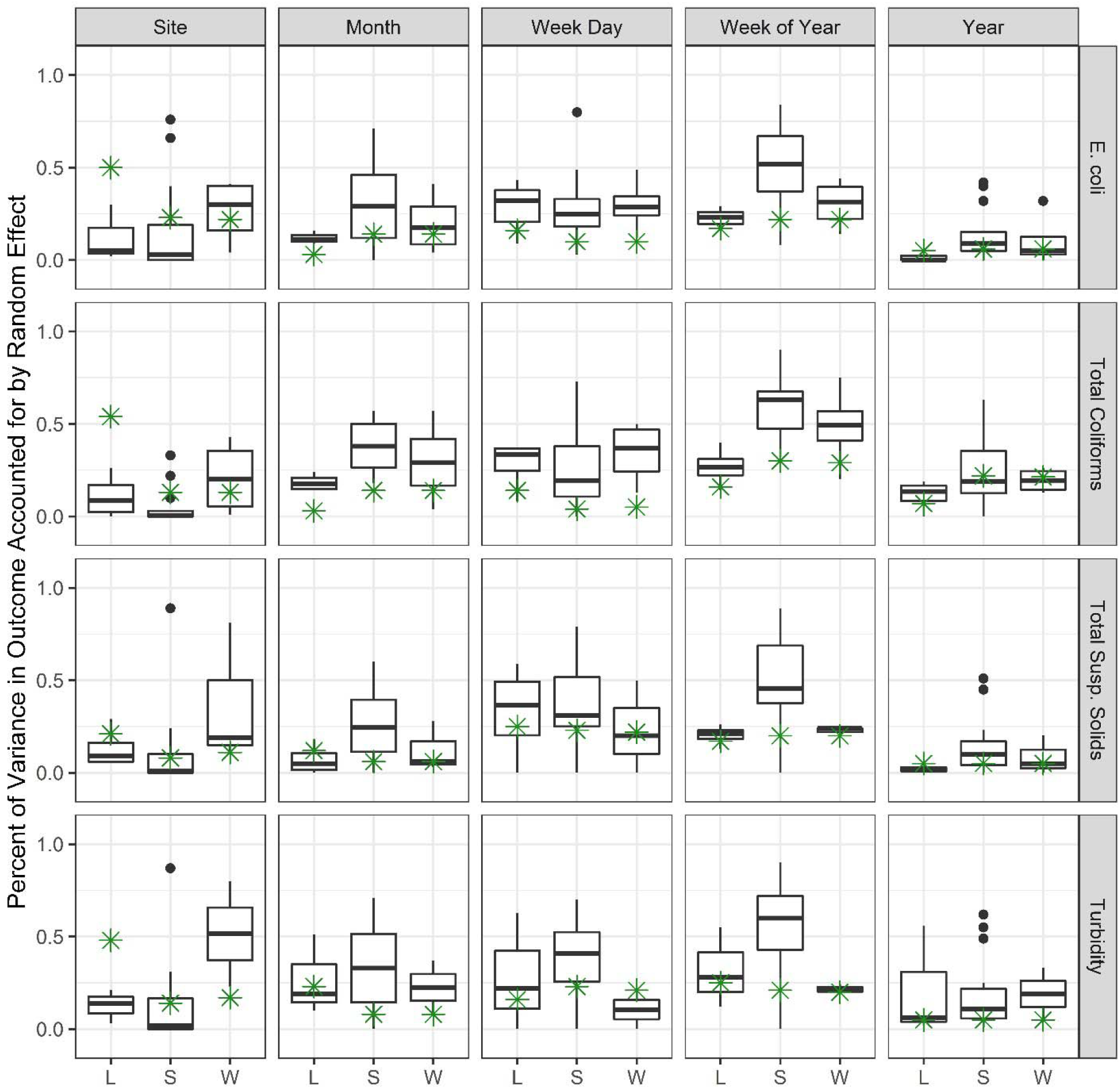
Variance in key water quality parameters in individual lakes (L), streams (S), and HUC-4/Finger Lake watersheds (W) accounted for when select spatial or temporal factors were included in random effects models (see Fig S4 for the full results). Results are only shown for lakes, streams, and watersheds with at least 20 samples. The variance in water quality for all lakes sampled, all streams, and all samples is represented by a green star in the corresponding column. Week of year = No. of weeks since Jan 1^st^.

**Figure 4:**
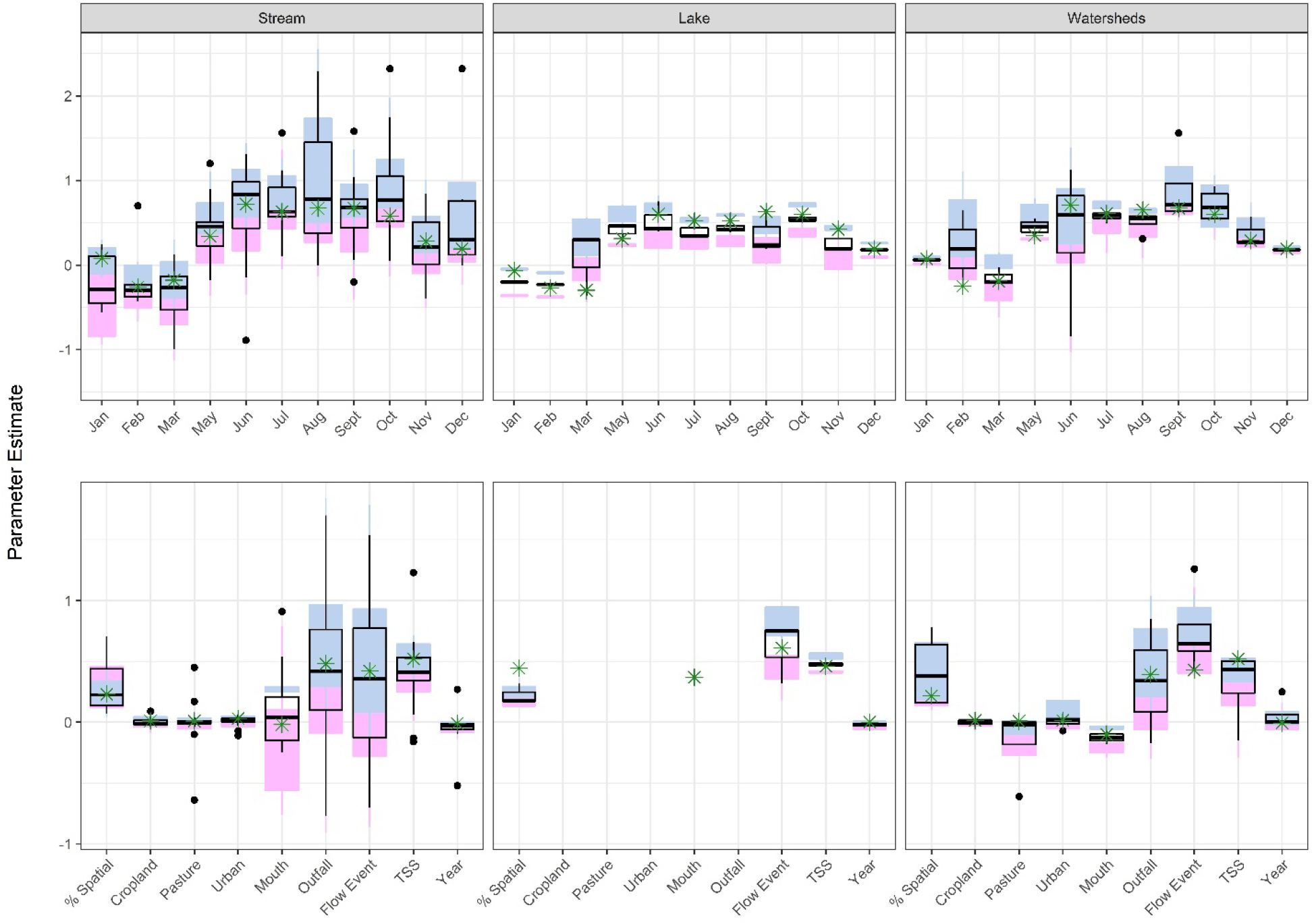
Results of a multivariable Bayesian models of coregionalization that assessed the association between environmental parameters and log10 *E. coli* levels. The 1^st^ and 3^rd^ quartiles, and median values for each estimate are shown. The boxplots show the distribution of the estimates for individual streams, lakes, and HUC-4/Finger Lake watersheds, while the estimate for models builts using the data from all lakes samples, all streams samples, and all samples is represented by a green star in the corresponding column. % spatial = percent of variance attributable to spatial effects. Urban, cropland, and pasture refer to the inverse-distance weighted proportion of the upstream watershed under these covers. TSS= Total suspended solids. The reference-level for mouth and outfall are these features being absent (as opposed to present), for flow event is baseflow (as opposed to stormflow), and for month is April.

### The strength and direction of associations between environmental parameters and log10 *E. coli* concentration were dependent on scale of analysis, water type, and waterway/watershed

Associations between environmental factors and log10 *E. coli* levels were assessed using multivariable Bayesian models of coregionalization (MCOR). Effect estimates in the MCOR can be interpreted as the change in *E. coli* levels (log10 CFU/100-mL) associated with a one-unit change in a continuous factor (such as log_10_ TSS) or with shifting from the reference-level to a different level of a categorical factor (e.g., from baseflow to stormflow conditions). While there was strong evidence of seasonal patterns in *E. coli* levels for both streams and lakes, the seasonal effect was more pronounced in the stream models compared to the lake models (Fig 4; Table S5). Regardless of waterway, watershed, water type, and scale of analysis, *E. coli* levels were highest in the summer and lowest in the winter (Fig 4; Table S5).

There also appeared to be a strong association between *E. coli* levels and TSS levels, and *E. coli* levels and flow conditions (Fig 4; Table S4). Specifically, *E. coli* levels were positively associated with TSS levels for all water types and scales of analysis as evidenced by the fact that the IQRs for the lake, stream, and watershed models did not include zero but did overlap each other and the effect estimates for the all-stream, all-lake and statewide models (Figure 4). According to the lake, watershed, all-lake, all-stream, and statewide models, *E. coli* levels were also higher under stormflow compared to baseflow conditions (Fig 4; Table S4), which may be evidence of event-scale effects (i.e., elevated *E. coli* levels during rain events). There was also evidence of an association between *E. coli* levels and flow conditions for most individual stream models, but the magnitude and direction of this effect were more variable for the stream models compared to the lake models or models with a larger scale of analysis (e.g., watershed, all-stream, and statewide models; Figure 4; Table S4).

While there was a small, non-negligible effect of land cover in the statewide models (effect estimates ranged between 0.01 and 0.03), some watershed and waterway models showed strong land-use effects (effect estimates ranged between −8.2 and 0.45; Figure 4; Table S4). Overall, the effect of land cover was dependent on scale of analysis and the waterway or watershed under consideration. Specifically, the absolute value of the effect estimates for land cover variables, particularly pasture, were generally but not always greater in the watershed compared to stream models. For example, a 1% increase in pasture cover was associated with a 0.00 (50% credible interval [CI]=0.00, 0.01) log10 CFU/100-mL increase in *E. coli* for the Seneca Lake watershed but a −0.61 (CI=−0.85, −0.40) log10 CFU/100-mL decrease for the Keuka Lake watershed. The effect estimate of land cover factors in many stream models was negligible, but there was a non-negligible effect for a subset of streams (e.g., Canoga Creek, Fall Creek, Kashong Creek, Keuka Outlet, Yawger Creek; Table S4). For example, a 1% increase in pasture cover was associated with a −0.10 (CI=−0.13, −0.06) decrease for Kashong Creek, a stream in the Seneca Lake watershed.

In contrast to land cover, there was a strong relationship between *E. coli* levels and outfalls across scales of analysis. However, the direction of this relationship varied between waterways (e.g., Big Stream versus Fall Creek) and watersheds (e.g., Cayuga Lake versus Seneca Lake), and between scales of analysis (e.g., statewide versus watershed models; Table S4). For example, when an outfall was present, the log10 CFUs/100-mL of *E. coli* increased by 0.39 (CI=0.30, 0.50), 0.85 (CI =0.66, 1.04), and 1.70 (IQR CI 1.57, 1.84) in the statewide, Seneca Lake watershed, and Keuka Outlet waterway models, respectively, but decreased by 0.17 (CI =−0.30, −0.06) and 0.77 (CI =−0.91, −0.62) in the Cayuga Lake watershed and Fall Creek waterway models, respectively (Table S4). The effect of sampling at the stream mouth (as opposed to upstream of the mouth) also varied with scale of analysis, the waterway/watershed under consideration, and water type. For example, *E. coli* levels were, on average, 0.91 (CI =0.79, 1.02) log10 CFU/100-mL higher at the Keuka Outlet mouth (compared to upstream sites) but 0.76 (CI =−0.99, −0.48) log10 CFU/100-mL lower at the Milliken Creek mouth (Table S4). Except for the all-lakes model, sampling at the stream mouth was associated with a decrease in *E. coli* levels at larger scales of analysis (e.g., statewide, all-stream, and watershed models); this may be due to the dilution effect as water moves from streams to stream mouths to lakes.

In models that included waterway type as a covariate, *E. coli* levels were substantially higher in streams and canals, and substantially lower in springs compared to lakes (Table S4). According to the statewide model and the Cayuga Lake watershed model, log10 *E. coli* levels were 0.43 CFU/100-mL (CI=0.29, 0.57), and 0.37 CFU/100-mL (CI=0.24, 0.49), respectively, higher in streams compared to lakes. Similarly, log10 *E. coli* levels were 0.27 CFU/100-mL (CI=0.10, 0.44), 0.19 CFU/100-mL (CI=0.03, 0.36), and 0.74 CFU/100-mL (CI=0.54, 0.95) higher in canals/flood control channels compared to lakes according to the statewide, all-lake, and Cayuga Lake watershed models, respectively. Lastly, in the statewide model, which included watershed as a covariate, *E. coli* levels were 0.69 log10 CFU/100-mL (CI =0.54, 0.84) higher in samples collected from the Keuka Lake Watershed, 0.25 log10 CFU/100-mL (CI =0.17, 0.33) higher in samples collected from the Seneca Lake Watershed, and 0.55 log10 CFU/100-mL (CI =0.73, 0.38) lower in samples collected from the Susquehanna River watershed compared to samples collected from the Cayuga Lake watershed, which was the reference-level.

### The distance at which microbial water quality became spatially independent increased as the scale of analysis increased but was also differed between models at the same scale

The effective range in the intercept-only MCOR models (the range at which *E. coli*, total coliform, TSS, or turbidity levels became spatially independent) varied substantially among models and scales of analysis (Table S2). According to the intercept-only MCOR, the effective range for *E. coli* was 778 m (N=385 samples; IQR=39, 1,511 m), 358 m (N=4,294 samples; IQR=259, 1,006 m), and 4 m (N=44 samples; IQR=3, 6 m) for the Seneca Lake, Cayuga Lake, and Susquehanna River watershed models (Table S2). The small effective range for the Susquehanna watershed may be an effect of the small number of Susquehanna watershed samples tested for *E. coli*; however, a similar number of Susquehanna watershed samples were =tested for TSS (N=47 samples; Median=59 m; IQR=41, 141 m) and turbidity (N=60 samples; Median=797 m; IQR=351, 2,799 m). The effective range for *E. coli* varied substantially between waterways as well. For instance, the effective range for *E. coli* in Cayuga Lake was 669 m (N=630 samples; IQR=498, 1,027 m) but was 117 m (N=65 samples; IQR=69, 279 m) in Keuka Lake and 6 m (N=84 samples; IQR = 4, 10 m) in Seneca Lake. While intensively sampled waterways with well-dispersed sampling sites, such as Cayuga Inlet (N=1,060 samples from 38 sites; Median = 96; IQR=57, 231 m) and Six Mile Creek (N=938 samples from 17 sites; Median=68 m; IQR=39, 120 m), often had small effective ranges for *E. coli*, several intensively sampled waterways with well-dispersed sampling sites had quite large effective ranges for *E. coli*, including Fall Creek (N=994 samples; Median = 721; IQR=464, 1,034) and Taughannock Creek (N=258 samples from 9 sites; Median=567 m; IQR=429, 843 m). Although the effective range for total coliforms, TSS, and turbidity similarly varied between waterways and watersheds (Table S2), the effective range for *E. coli* did not appear related to the effective range for total coliforms, TSS, or turbidity in a given waterway/watershed. For instance, the effective range for total coliforms, TSS and turbidity in Cayuga Inlet were 162 m (N=924 samples; IQR=110, 356 m), 7,754 m (N=989 samples; IQR=429, 843 m), and 3,931 m (N=1,053 samples; IQR=3,658, 3,989 m), respectively (Table S2). Despite this variability in effective range for a given parameter between waterways and watersheds, and between parameters for a given waterway/watershed it is important to note that for most waterways the effective range was consistently low (i.e., measured in the 10s or 100s of m as opposed to 1000s of m). Indeed, 64% (14/22) of *E. coli* stream models, 58% (11/19) of total coliform stream models, 57% (13/23) of TSS stream models, and 67% (14/21) of turbidity stream models had an effective range less than 100 m, while approx. 40% had an effective range <10 m, and 14% had an effective range >500 m. It is also important to note that in both the intercept-only and multivariable MCOR, effective range generally increased as spatial scale increased and, for total coliforms, TSS and turbidity, was generally larger for the lake as opposed to the stream models. For example, the median effective range for all *E. coli* intercept-only watershed MCOR was approx. 4.5 times greater (261 m) than the median effective range for all 22 *E. coli* stream MCOR (58 m).

## Discussion

### The scale of analysis strongly affected the observed sources of variance

The variance attributable to any given factor in the waterway and watershed models ranged widely with large differences among waterways. This finding, that sources of variation in microbial water quality were waterway-specific is supported by previous studies that reported waterway-specific findings and attributed this to the heterogeneity inherent to freshwater environments (e.g., (13, 26, 30, 53)). Despite these differences, as spatial scale increased (from the waterway to watershed to statewide models), the variance attributable to spatial sources generally increased and the variance attributable to temporal sources generally decreased in the present study. This finding is consistent with past studies that associated FIB levels with stream order/watershed size and concluded that observations at coarser scales could reveal patterns (and drivers of these patterns) not detectable at finer scales (34, 54–56). Conversely, the dilution of local effects when data are aggregated to coarser scales is well-established (55, 57). For example, a study of spatial autocorrelation in *E. coli* levels in the Norwalk River, CT, USA found that signals from point sources were rapidly diluted as water moved downstream, with *E. coli* levels being ≥90% lower 1,000 m downstream of a potential source than at the source (36). Based on the effective ranges (the distance at which *E. coli* levels became spatially independent) in the present study, the distance at which dilution occurred varied substantially between waterways and across scales of analysis (again increasing with increasing scale), but was consistently below the 1,000 m reported previously (36). Overall, these findings highlight the importance of considering scale when designing monitoring programs. For example, the Norwalk River study and this study, both suggest that monitoring programs designed to identify point sources of contamination should place sampling sites ≤ 1,000 m apart. However, such intensive sampling may be cost-prohibitive and should be balanced against affordability. Composite sampling may provide an alternative way to account for spatial variability while reducing the cost of testing multiple samples (58–62). Indeed, multiple studies determined that compositing samples did not produce statistically different FIB counts or result in different management decisions compared to analyzing each sample individually (58, 59, 61, 62). Adapting the intensity of sampling based on the probability of contamination (more sites near point sources and fewer sites, spaced farther apart further from the sources) might also work.

In addition to considering the scale of analysis, monitoring programs should also be adapted to different types of waterways (e.g., lakes, streams) and sampling site (e.g., stream mouth, areas near outfalls). In the present study, more variance in microbial water quality was attributable to spatial factors in the lake compared to the streams models, possibly because the sampled lakes (e.g., Cayuga Lake, Seneca Lake) were considerably larger than the drainage basins of the sampled streams. However, more variance was attributable to spatial sources in the all-lake model than the all-stream model even though the all-stream model represents a larger area than the all-lake model, which may indicate that this finding is not solely an artifact of differences in drainage basin size. Indeed, these findings are supported by Partyka et al. (53), who compared FIB levels in six California reservoirs along a transect from the reservoirs’ inflows to their outflows and found significant differences in *E. coli* coliform levels between inflow, outflow, and reservoir samples. These findings also make sense when microbial fate and transport and waterway dynamics are considered. Drivers of microbial contamination and factors that facilitate microbial survival are likely to differ between standing bodies of water (e.g., lakes) and flowing waterways (e.g., streams). For example, while sediments are important in-channel stores for microbial contaminants (63, 64), the influence of sediments on the water column is affected by waterway-specific characteristics, including particle size, flow rate, and depth (65, 66). Thus, the influence of sediments is likely less in the deeper, slower-moving lakes than the shallower, fast-flowing streams sampled here. For example, where Taughannock Creek enters Cayuga Lake depth changes from approx. 0.30-1.2 m at the stream-mouth to approx. 3-15 m in the near-shore area, and approx. 100 m at the lake center (based on topographic maps, usa.fishermap.org/depth-map/cayuga-lake-ny/). As a result, resuspension of sediments can be expected to have a greater effect on microbial and sediment levels in Taughannock Creek compared to Cayuga Lake purely due to depth and dilution. Overall, our study suggests that different monitoring strategies may be needed to fully capture spatiotemporal variation and assess microbial water quality for different types of surface waterways, particularly if monitoring will be used to guide decisions with public health implications. Our results also suggest that one-size-fits-all regulatory standards may not be appropriate. Instead, standards tailored to specific water types may be appropriate; however, this requires further investigation to assess the efficacy of water type-specific standards.

### Intra-annual variability accounted for more of the temporal signals observed here than interannual variability

This study assessed variation during events (baseflow versus stormflow sampling), over a year (represented by week-of-the-year, month, and season), and between years. The descriptive data visualizations and the random effects models showed limited interannual but considerable interannual variation in both FIB and sediment levels. Moreover, substantial variance was attributable to seasonal and event-scale variables. Indeed, in the multivariable lake, stream, and watershed models, the largest magnitudes of effect were attributed to month and flow event, while the effect of year was negligible. Past studies also reported strong seasonal (39, 67, 68), event-scale (39), and weekly patterns in microbial water quality. For instance, a review on sources of temporal variability in microbial water quality reported that event-scale variability was the greatest source with FIB levels changing significantly in response to rain events (39). Since seasonal, event-scale, and weekly patterns were observed by studies conducted in different regions and waterways using different sampling designs, microbial targets, and laboratory methods suggests that the temporal patterns observed here are reproducible.

Because of the dominance of event-scale and seasonal variation in FIB levels, multiple studies have suggested that compliance with the proposed *E. coli*-based FDA agricultural water quality standard (which is calculated using just 20 samples over a 2 to 4 year period) is highly dependent on when the =samples were collected (13, 25, 28). Thus, compliance with the proposed standard may not be indicative of water quality at time of water use, and compliance with the standard may not be an appropriate basis for guiding risk management decisions (e.g., when and where water should be used for irrigation or recreation). Instead, standards and associated monitoring programs, that account for seasonal and event-scale variability may be more useful for guiding these decisions. For instance, sampling could be targeted to conditions known to be associated with increased FIB levels (e.g., summer months, stormflow conditions) to provide a more conservative estimate of microbial contamination risks. Monitoring programs should also consider when and how the water is used. For example, agricultural water use in the Northeastern US, where the present study was conducted, is likely to peak in summer and fall. Monitoring programs that more intensely sample during summer and fall due to seasonal patterns in FIB levels (i.e., increased levels in these seasons), would also be targetting periods with an increased chance of human exposure (e.g., through irrigation). In considering how to adapt standards and monitoring programs, it is also important to note that, a single surface water source may be used in multiple ways that are weather-independent and unrelated to crop water needs, such as pesticide application, dust abatement, frost protection, and fertigation.

Our results also suggest an interesting connection to recreational activities since a non-negligible amount of variance was attributable to day of the week across all scales of analysis. FIB levels tended to be lower during the work week and higher over the weekend with the highest levels on Friday and in the Spring. Previous studies concluded that recreational activity (e.g., camping, hiking, swimming) on the weekends can affect the microbial quality of downstream surface water sources (69–73). Two studies conducted in Washington State found that fecal coliform and fecal streptococci concentrations in surface water were higher on weekends (when approx. 90% of recreational activities occurred) compared to weekdays (69, 70). All waterways sampled in the study reported here are in Upstate New York, which has large tourism and outdoor recreation industries. The authors personally observed campgrounds, hiking pull-offs, swimming areas, public fishing sites, and vacation rentals upstream of or near many of the sampling sites. However, weekly patterns in other human activities, such as agriculture, could also explain the variance attributable to weekday in the present study.

### While the effect of land cover on *E. coli* levels was highly dependent on scale of analysis, the magnitude of the land cover effect was consistently smaller than the effect of outfall presence

The effect of land cover was largely dependent on scale of analysis and the waterway or watershed under consideration. The effect estimates for land cover factors in waterway models were mostly negligible, although there was a non-negligible effect for a subset of streams. Conversely, the magnitude of effect for wastewater outfall presence and site type (i.e., if the sampling site was a stream mouth or not) was consistently larger than the magnitude of effect for land cover factors. Overall, this is consistent with past studies, which also found that the effect of land cover was dependent on the spatial scale considered (37, 38, 74). One study of 40 Canadian rivers conducted to determine the scale at which land cover was associated with *E. coli* levels found that land cover and *E. coli* levels were most strongly associated at smaller spatial scales, between 5 and 10 km (38). A New Zealand study found that the relationship between land cover and FIB levels was dependent on stream size and the area considered when calculating land cover with local land cover being more strongly associated with FIB levels in small streams and watershed-scale land cover having a stronger influence on FIB levels in large streams (37). Based on these findings, the New Zealand study concluded that FIB monitoring programs need to be scale-sensitive; a conclusion supported by our findings and the Canadian study (38).

The effect estimate for outfall presence (a point feature) was greater than the effect estimates for all three land-use factors (cropland, pasture, and forest-wetland cover) at all scales of analysis considered in the present study. Other studies also found that impaired water quality was associated with wastewater and other outfalls presence (75–77), including a study that sampled several of the streams sampled in our study (12). A survey of the Seine watershed in France found that microbial inputs from point sources (e.g., wastewater outfalls) were substantially greater than non-point source inputs associated with land use patterns under baseflow conditions, and recommended prioritization of remediation efforts to mitigate microbial and fecal inputs from point sources as opposed to overland run-off from non-point sources (77). The present study largely supports this conclusion. However, the direction of the relationship between point sources and *E. coli* levels was negative for a minority of models, possibly because outfall presence was confounded with sampling at the stream mouth or impoundments and pools. The aforementioned dilution effect may have obfuscated the point source effect at such sites (39). Indeed, in the present study sampling at a stream mouth or pond was associated with reduced *E. coli* levels compared to streams.

Throughout the 2010s, produce industry stakeholders have highlighted concerns about the agricultural water standard proposed as part of the Produce Safety Rule (29). One concern repeatedly raised at topical summits, conferences, and other venues was the lack of guidance on sampling, including how close together irrigation pumps that drew from the same source needed to be considered the same source for monitoring and compliance purposes (29). Thus, in considering the implications of our findings for monitoring programs, it is important to consider the distance at which microbial water quality became spatially independent. Although this distance increased as scale of analysis increased in the present study, it was variable and consistently low for most waterways (i.e., measurable in the 10s of m as opposed to 1000s of m). This indicates that FIB levels in samples collected relatively close together (e.g., < 1 km), would be spatially independent. Therefore, sampling at indiviudal irrigation intakes, as opposed to treating a single sampling site as representative of water quality at multiple intakes, may be needed to ensure compliance with the proposed standard. Thus, the findings reported here not only highlight the importance of scale when designing monitoring studies, they also bring into question the utility of one-size-fits all standards and instead suggest that standards tailored to specific water types may be appropriate. However, this requires further investigation in follow-on studies and/or meta-analyses specifically aimed at assessing the efficacy of water type-specific standards.

## Supplemental Materials

**Table S1:**
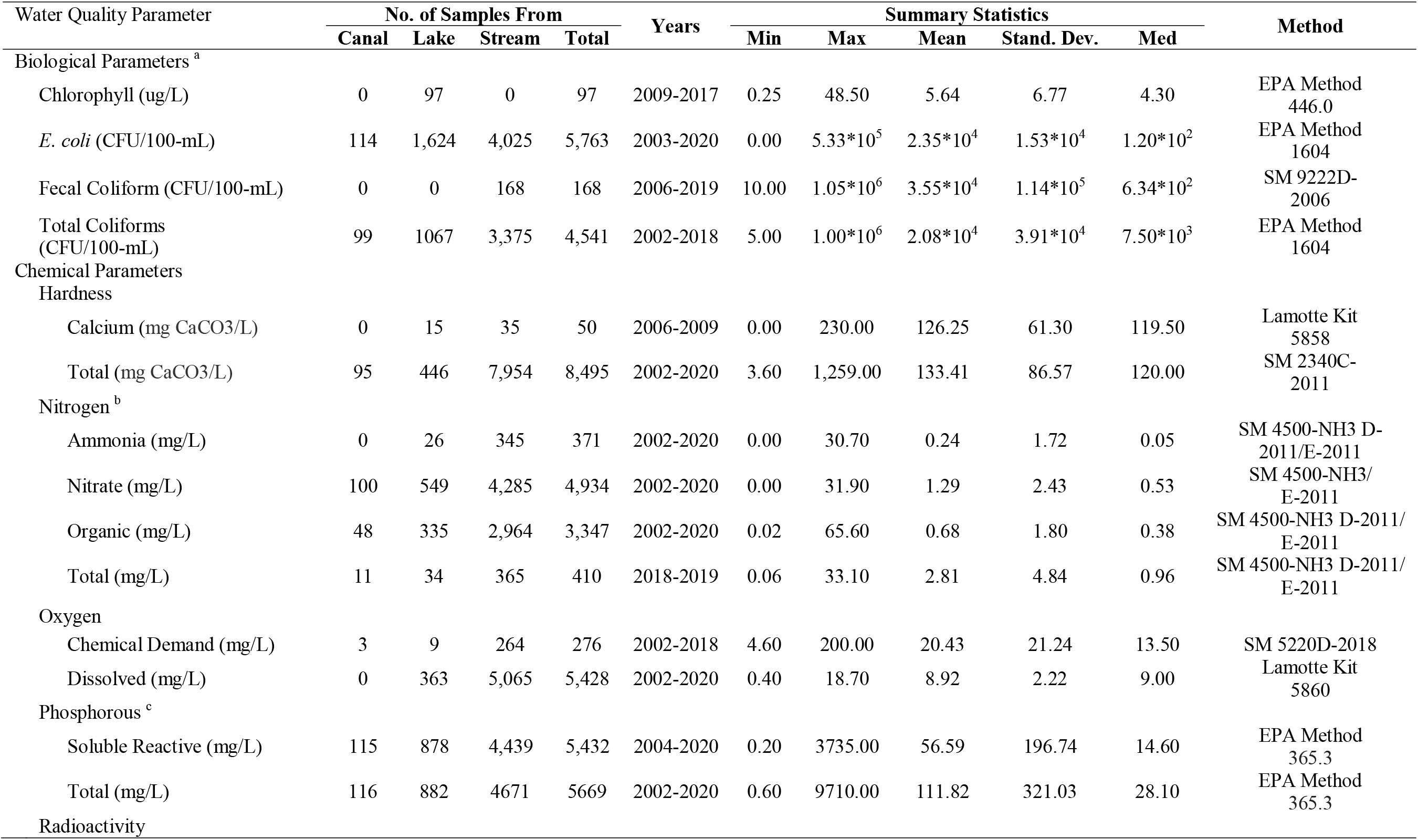
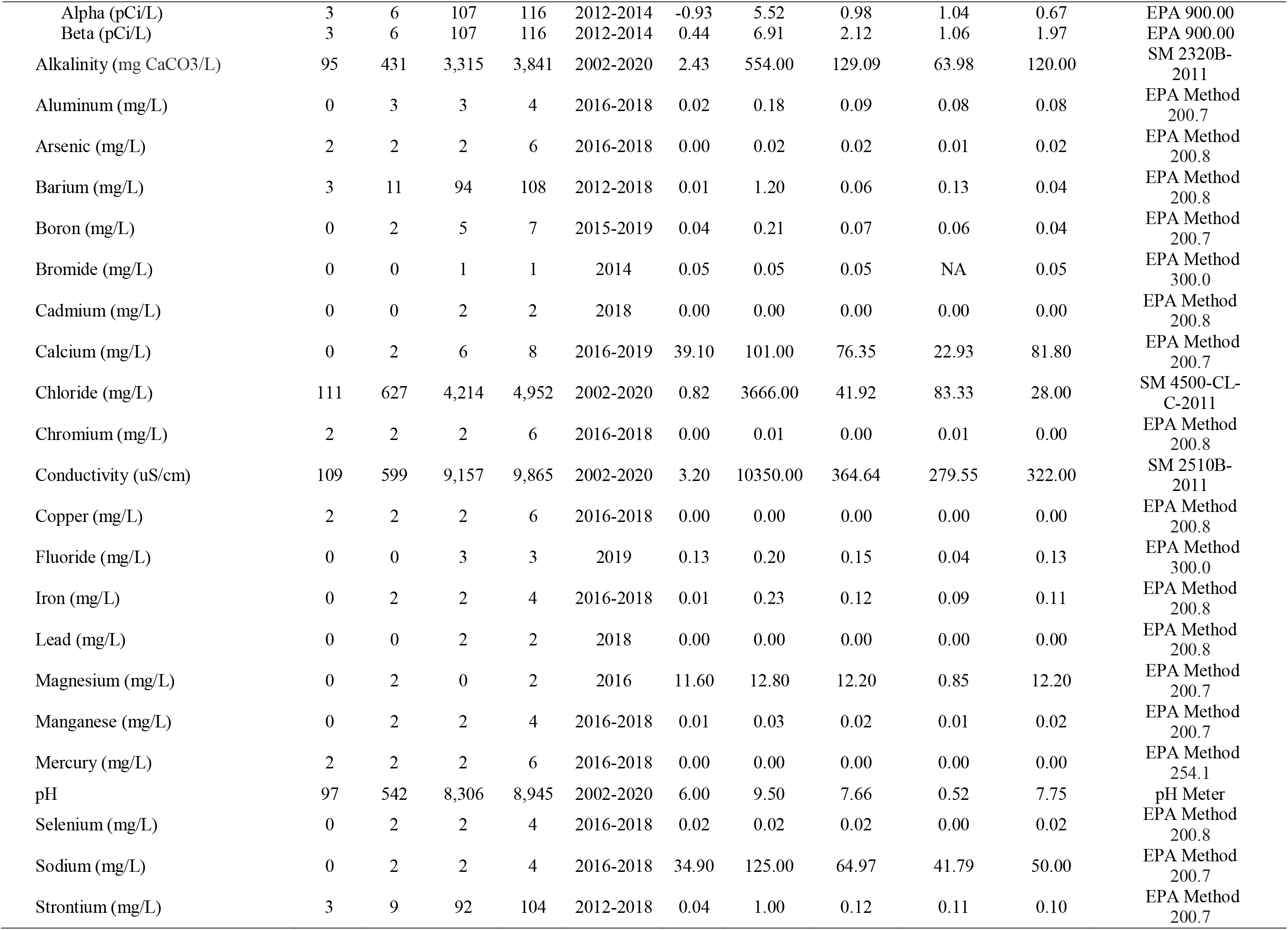
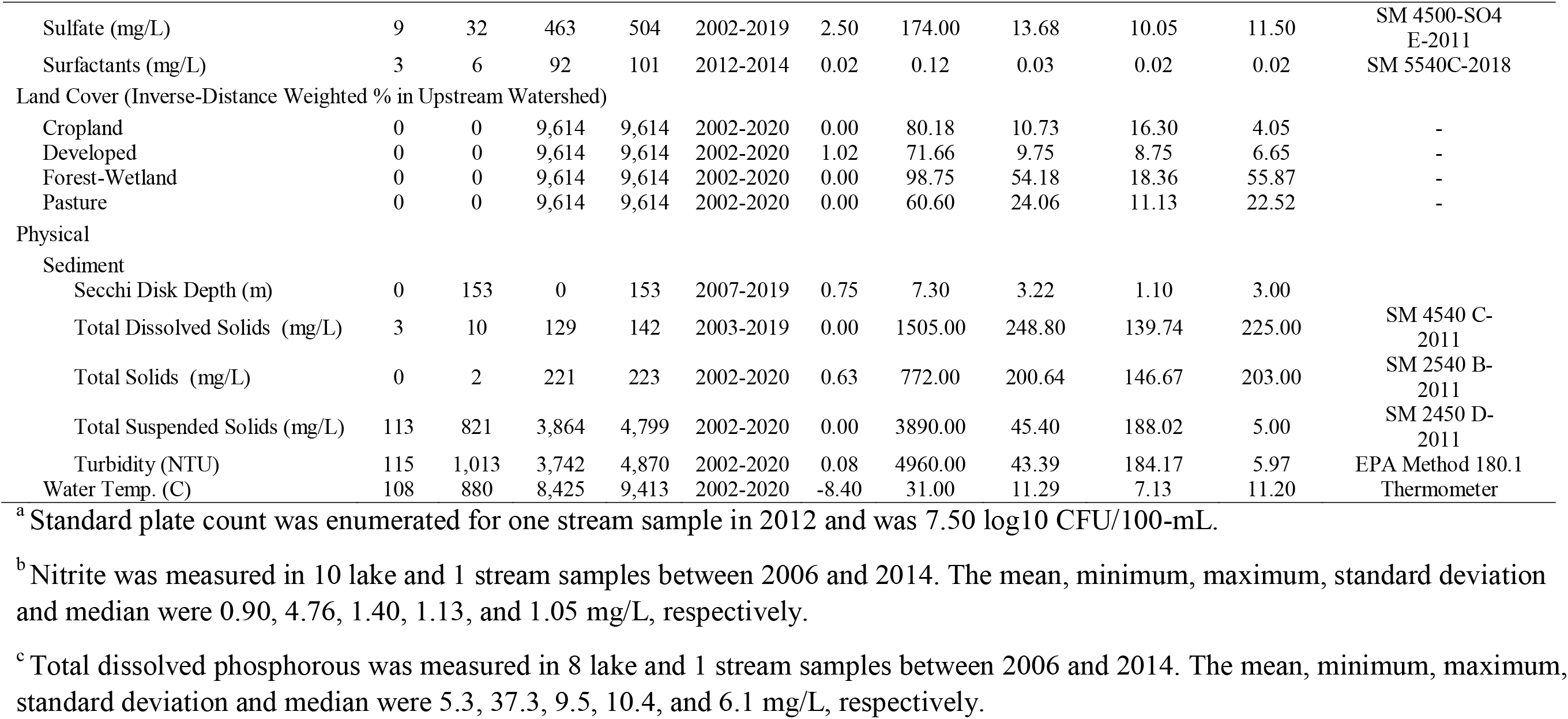
Summary statistics for biological, chemical, and physical water quality parameters in Upstate New York surface water sources sampled between 2000 and 2020; five land-use parameters are also summarized. The laboratory method used is cited in the final column.

**Table S2:**
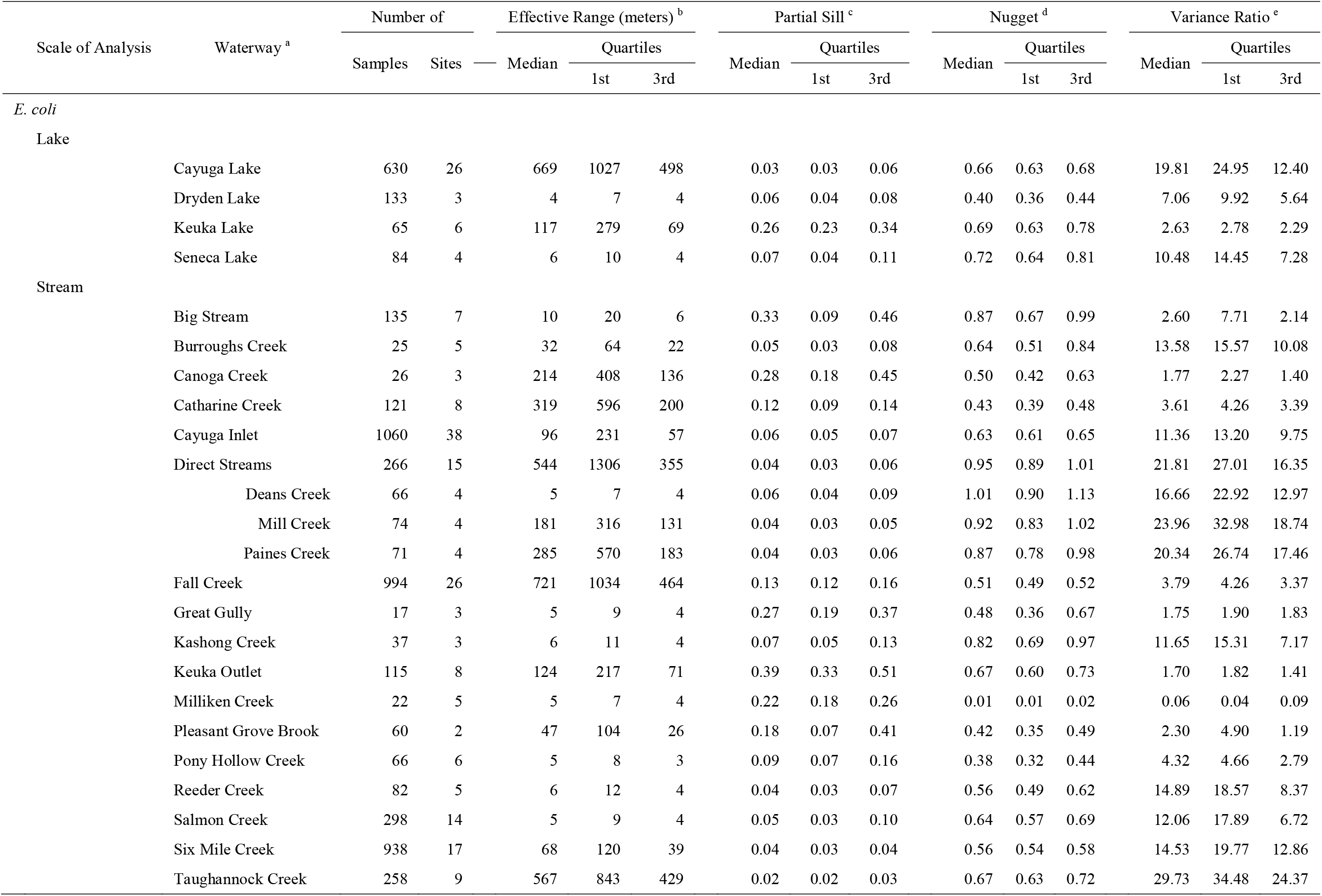
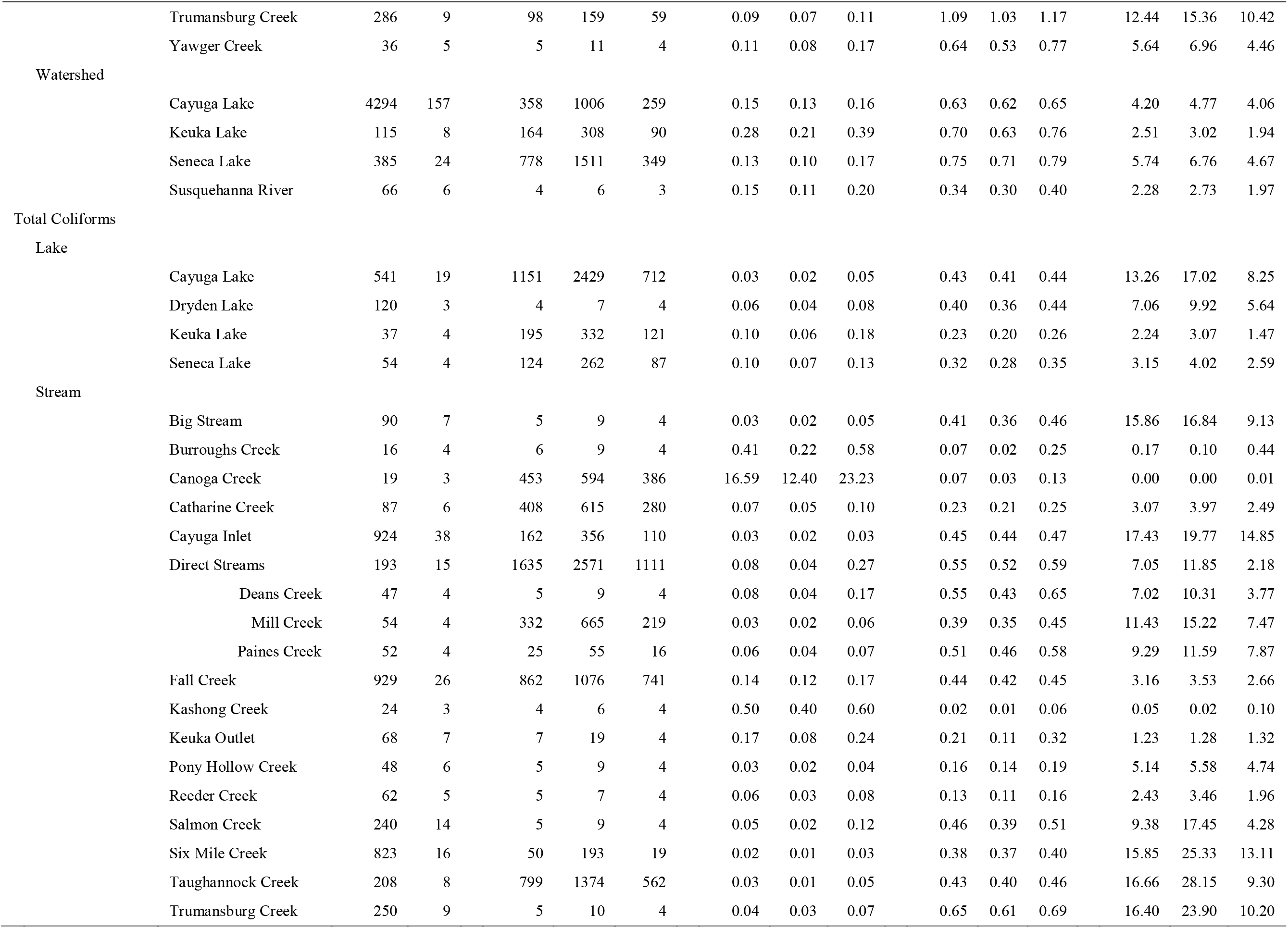
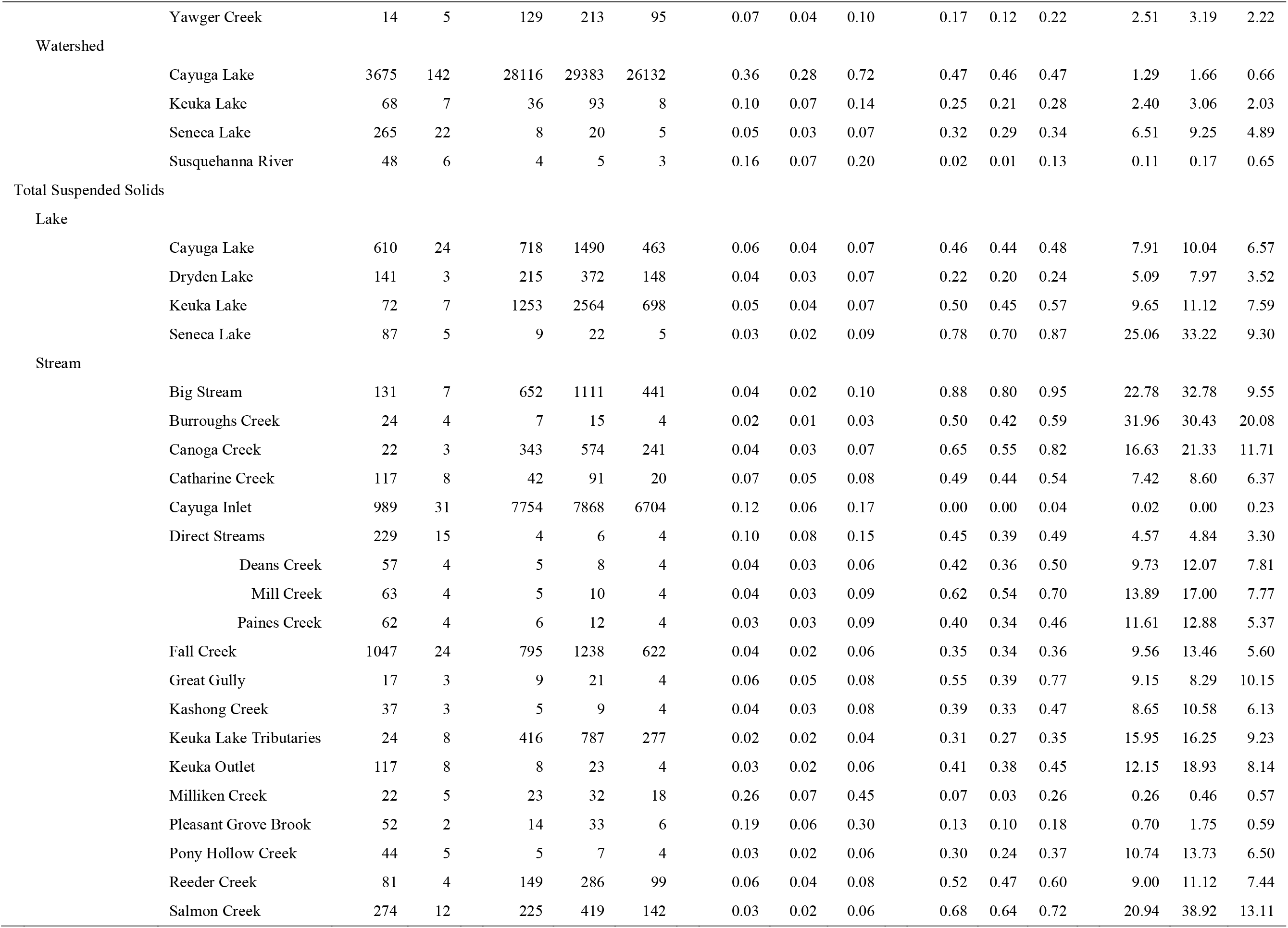
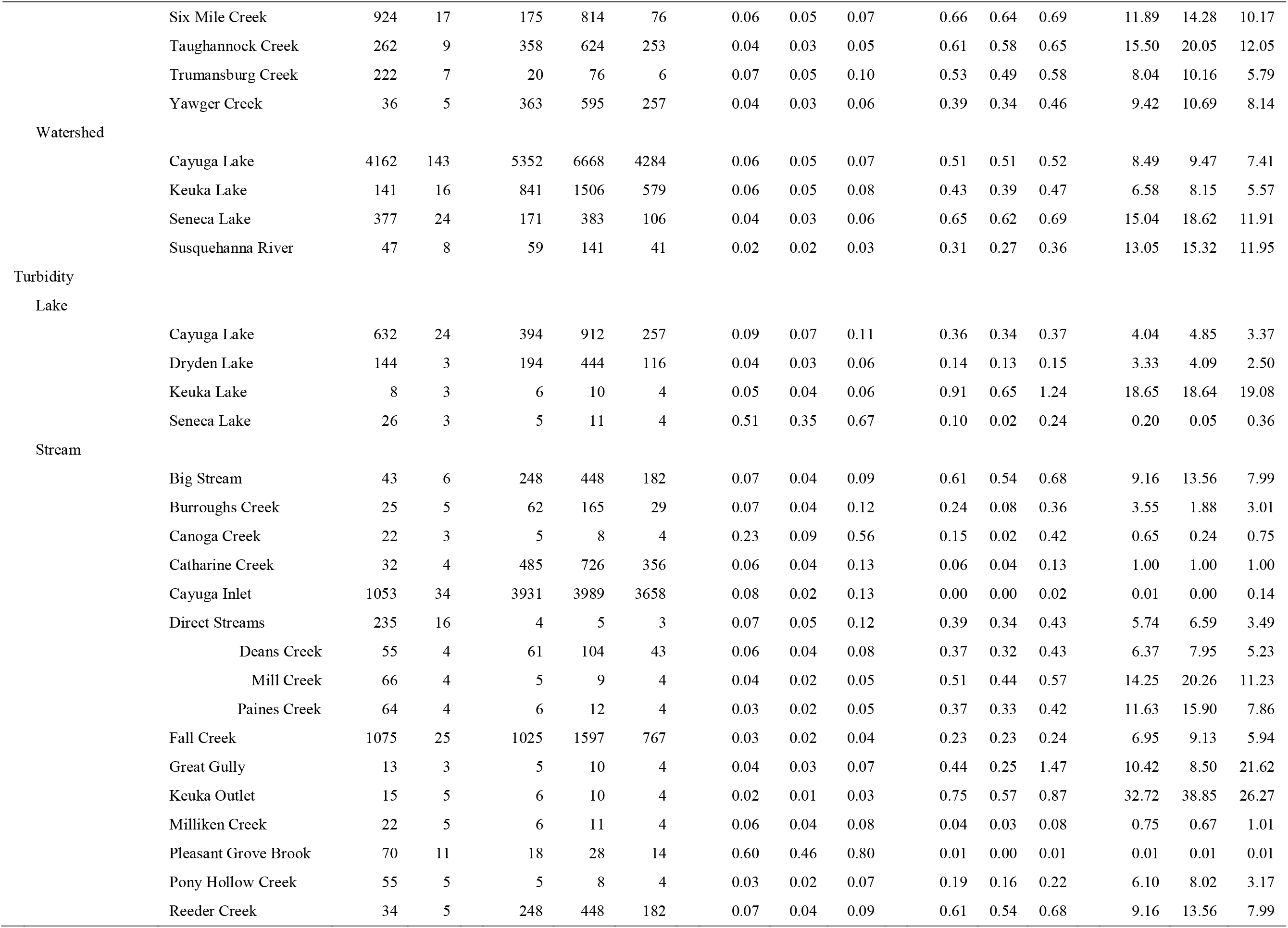
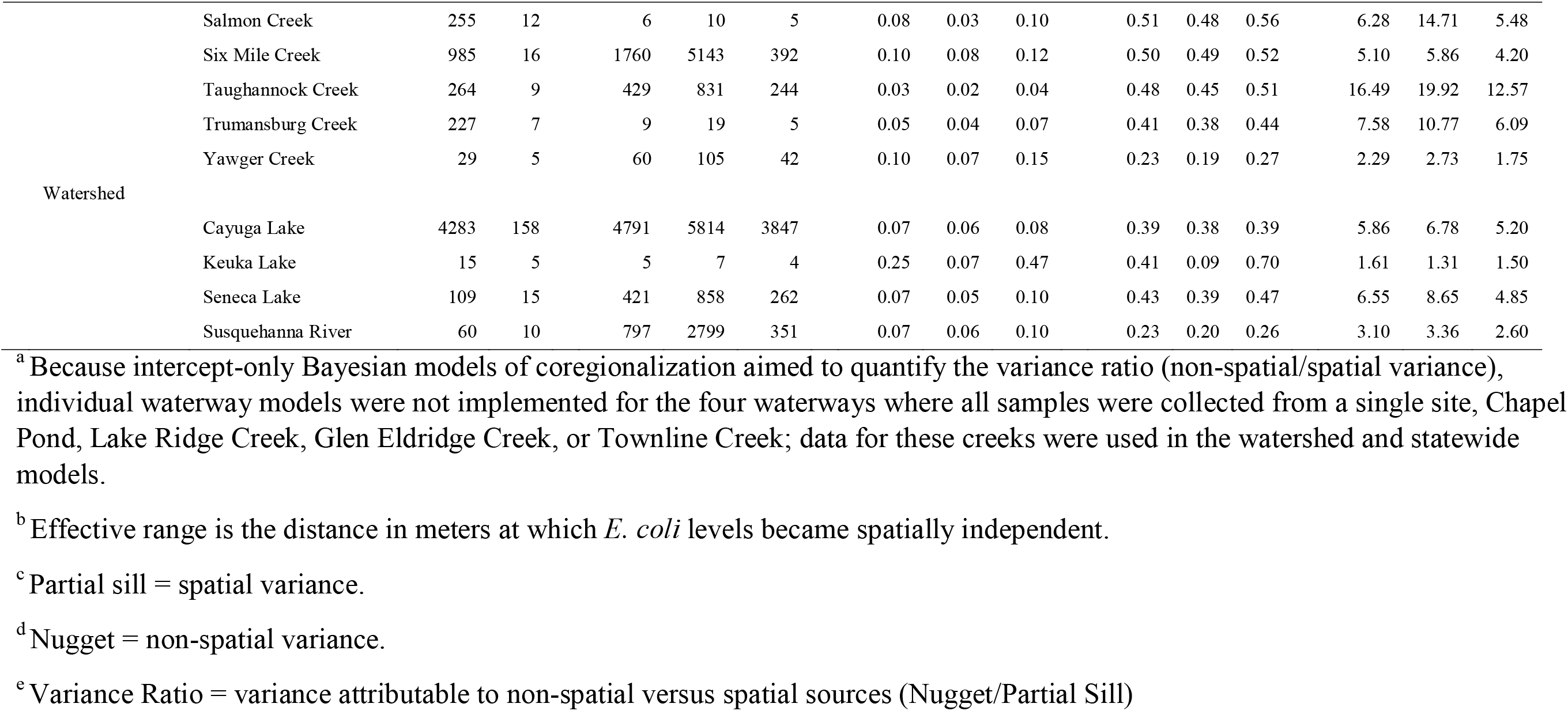
Results from the intercept-only Bayesian models of coregionalization.

**Table S3:**
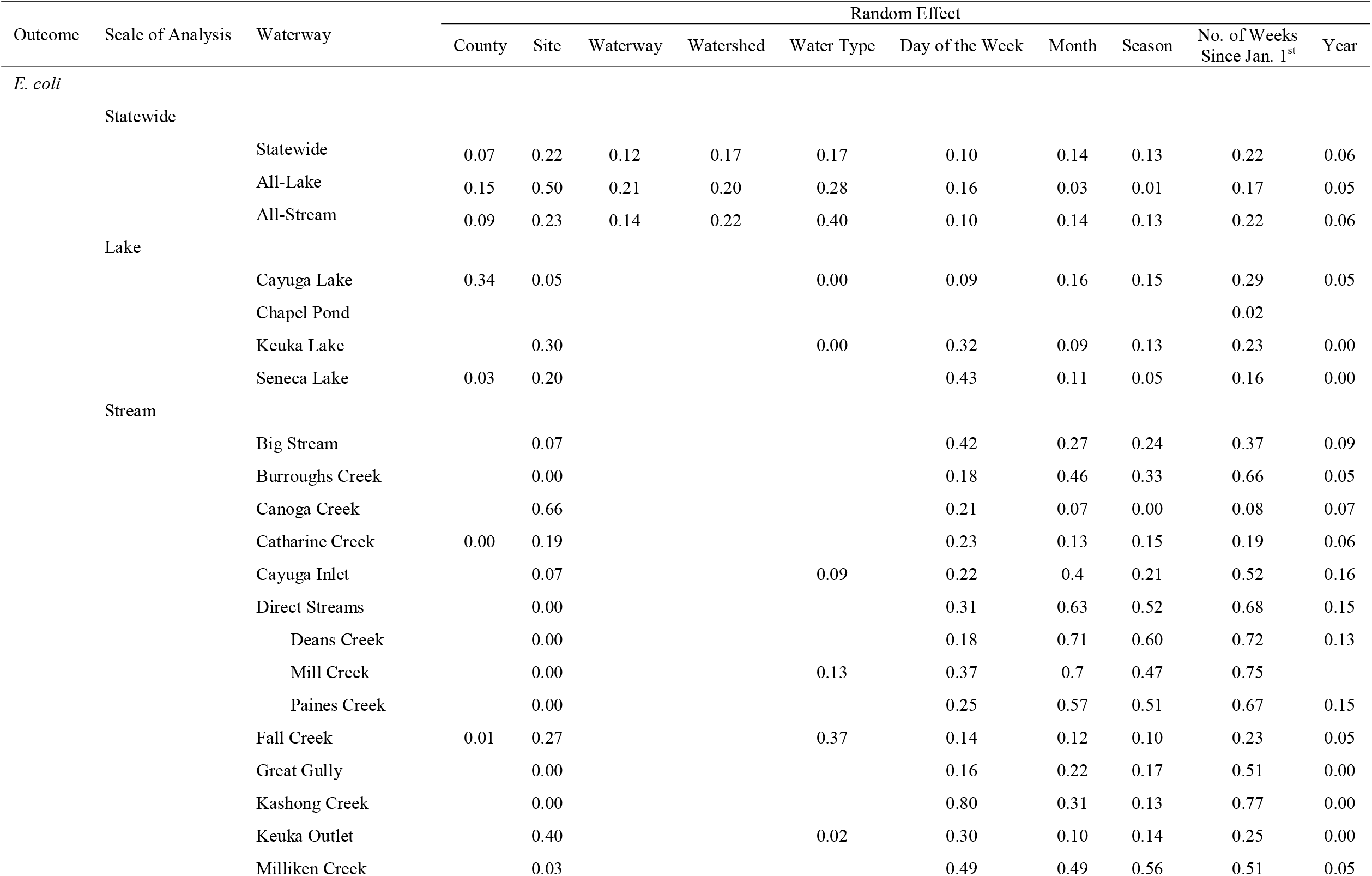
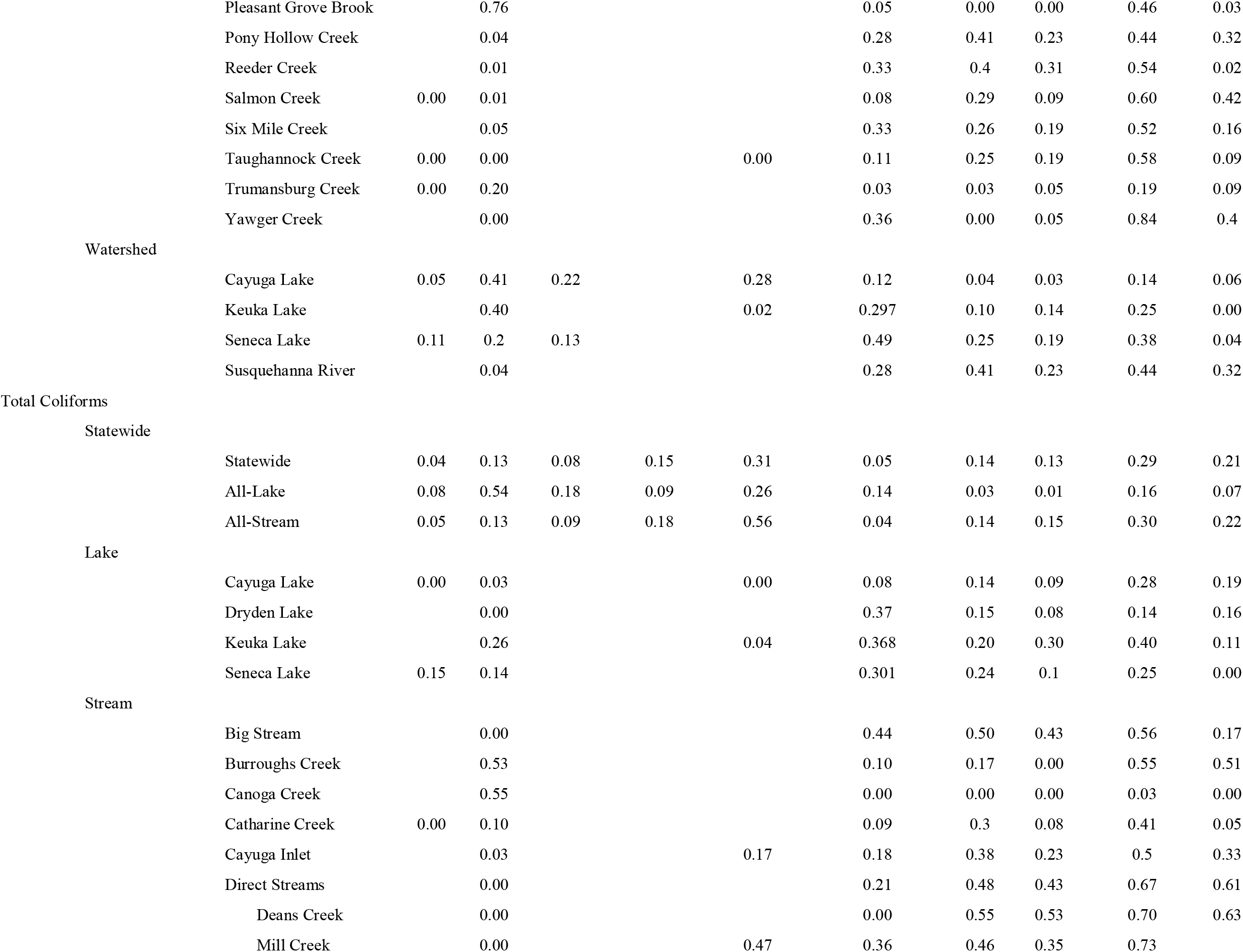
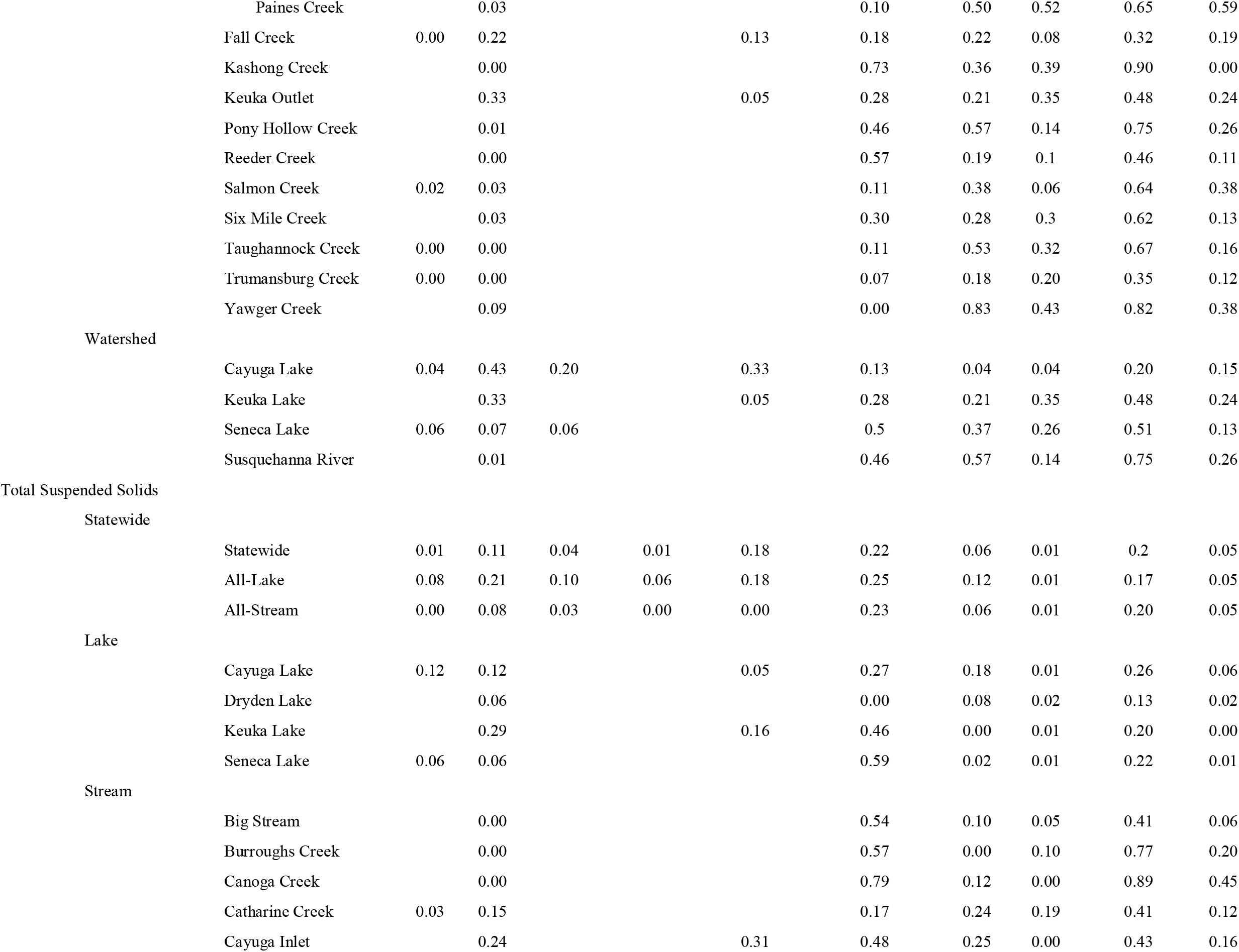
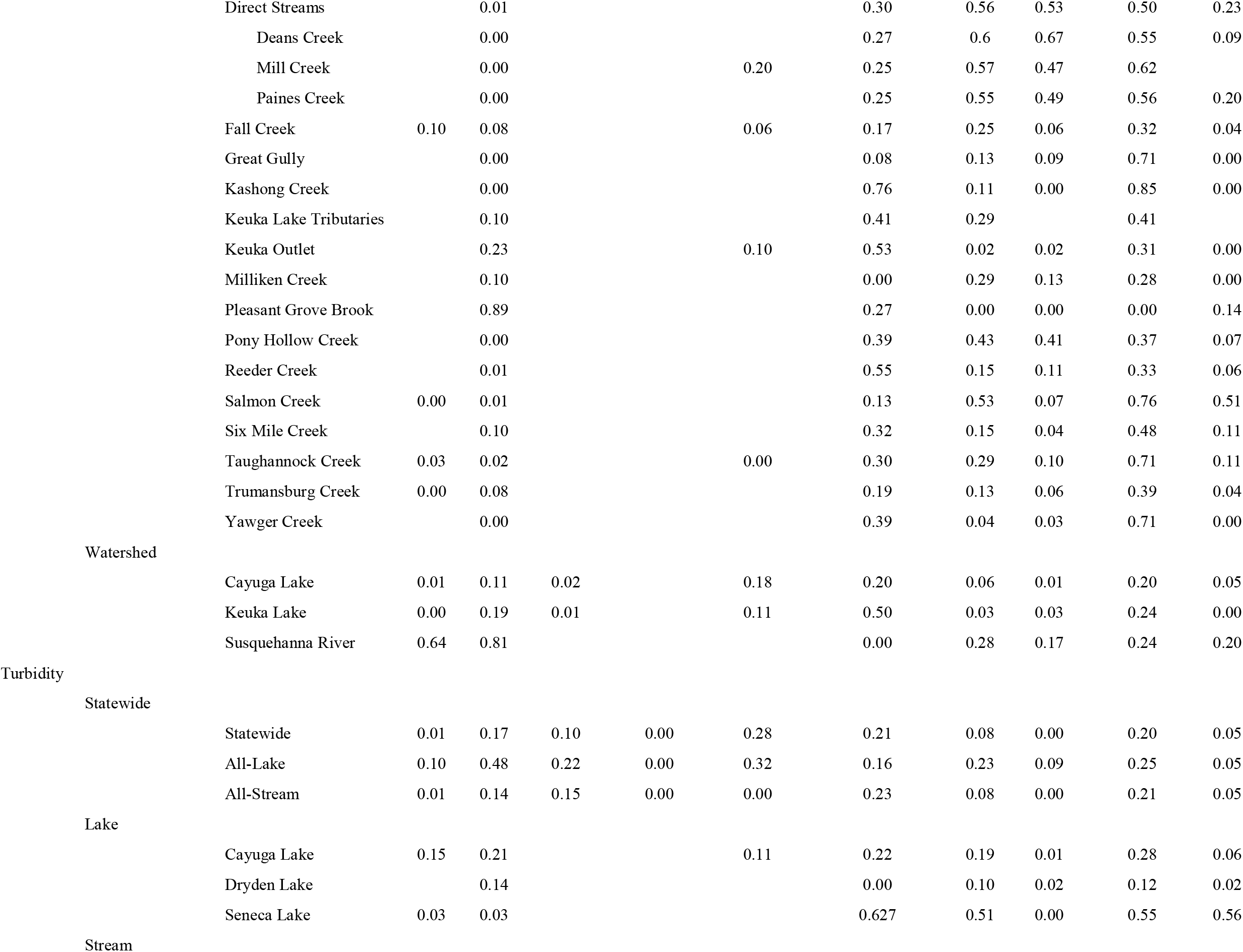
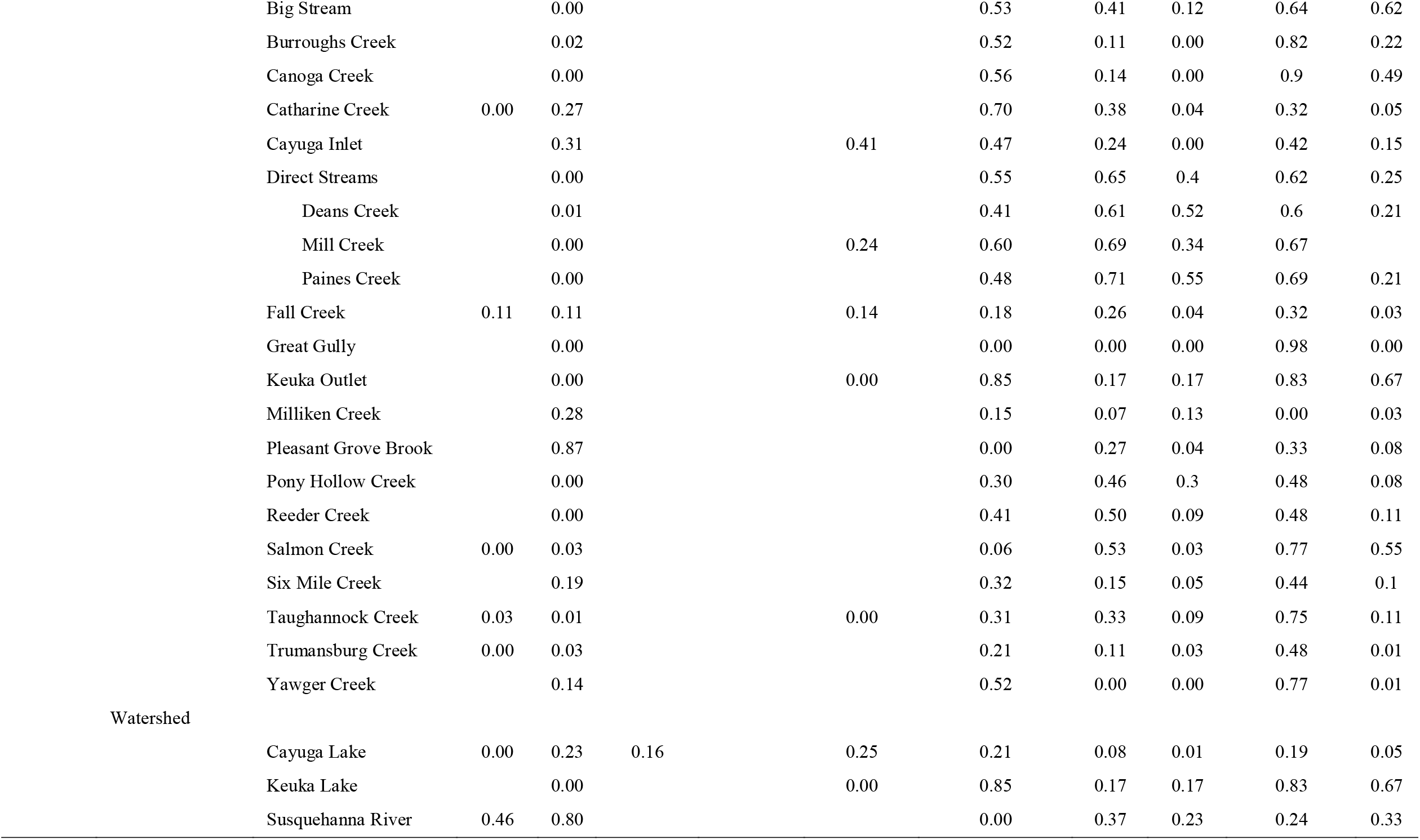
Results from the random effects models; blank cells indicate that for the given combination of parameter, scale of analysis, and waterway that the model could not be implemented for the random effect in the column. For instance, waterway could not be included as a random effect for the waterway models. Similarly, all of the sites along Big Stream were in a single county so county could not be included as a random effect for the Big Stream models.

**Table S4:**
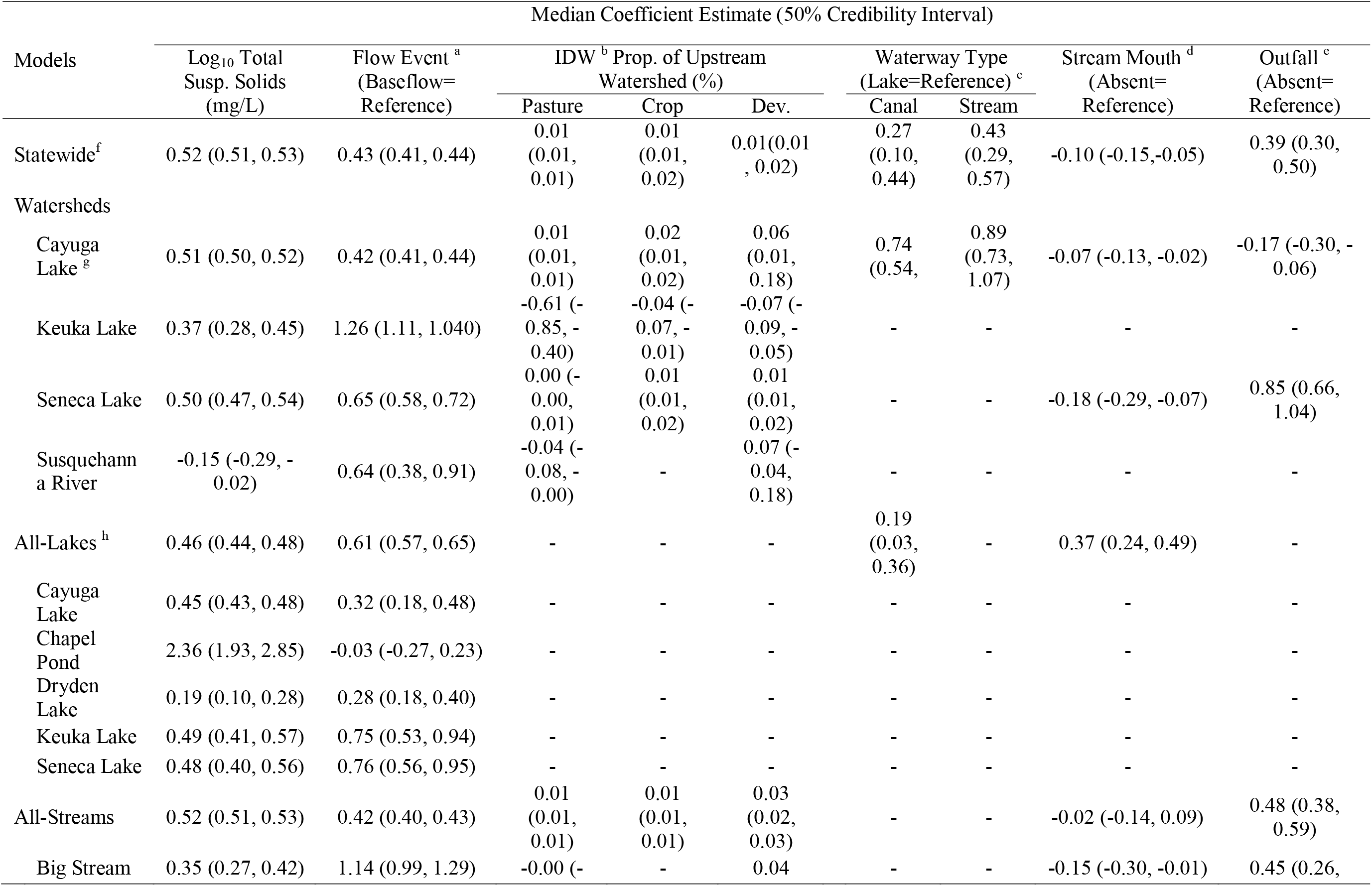
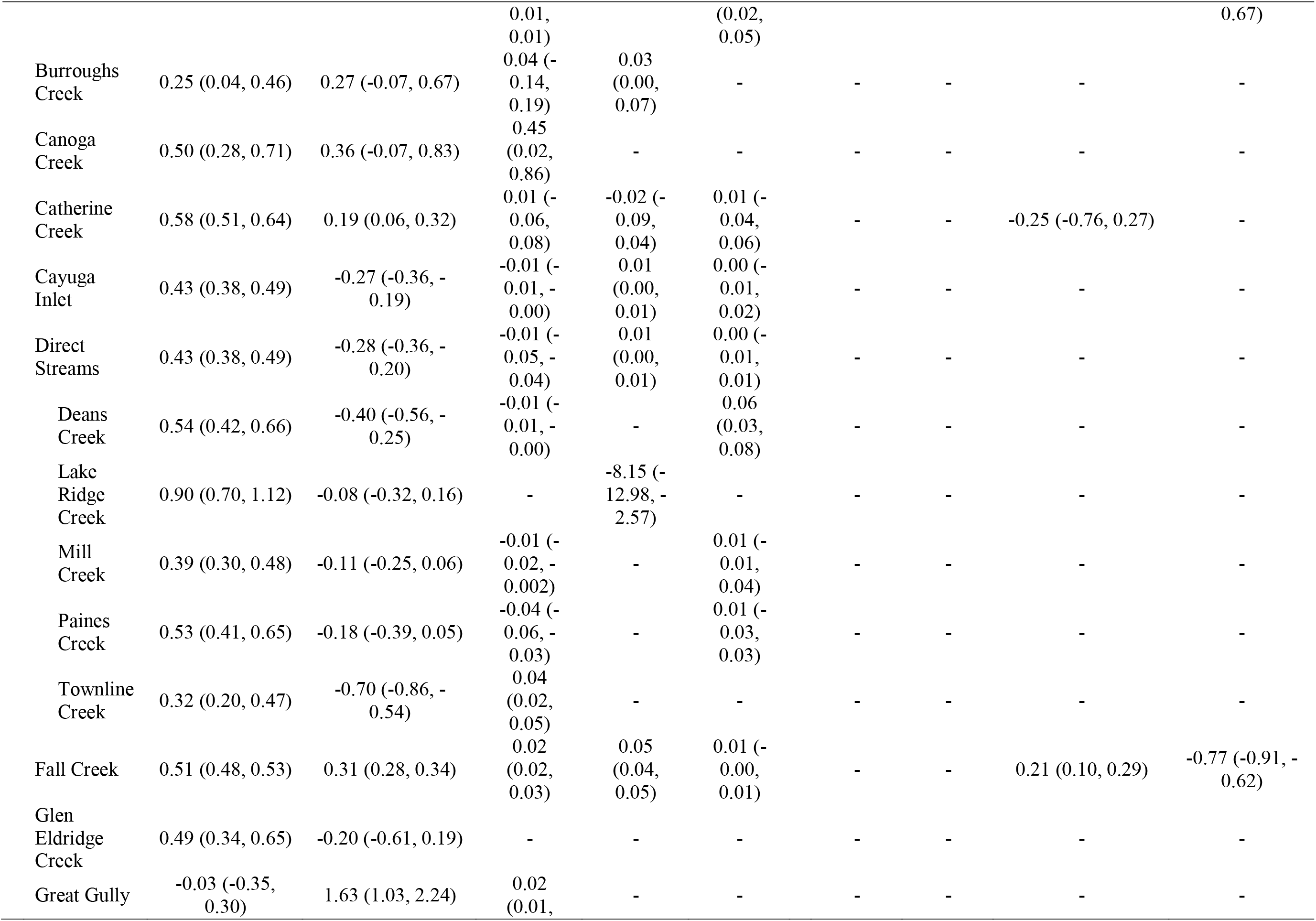
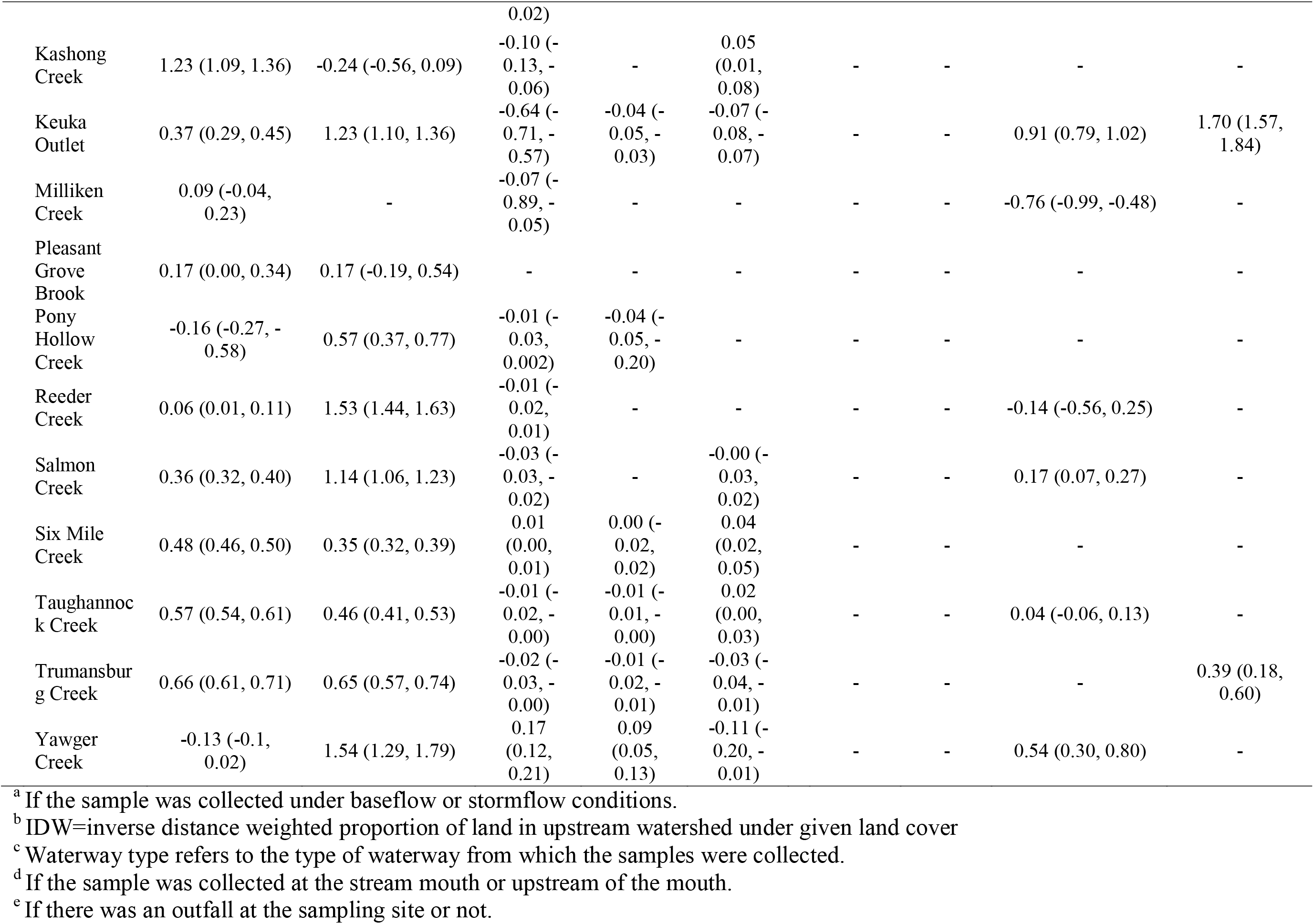
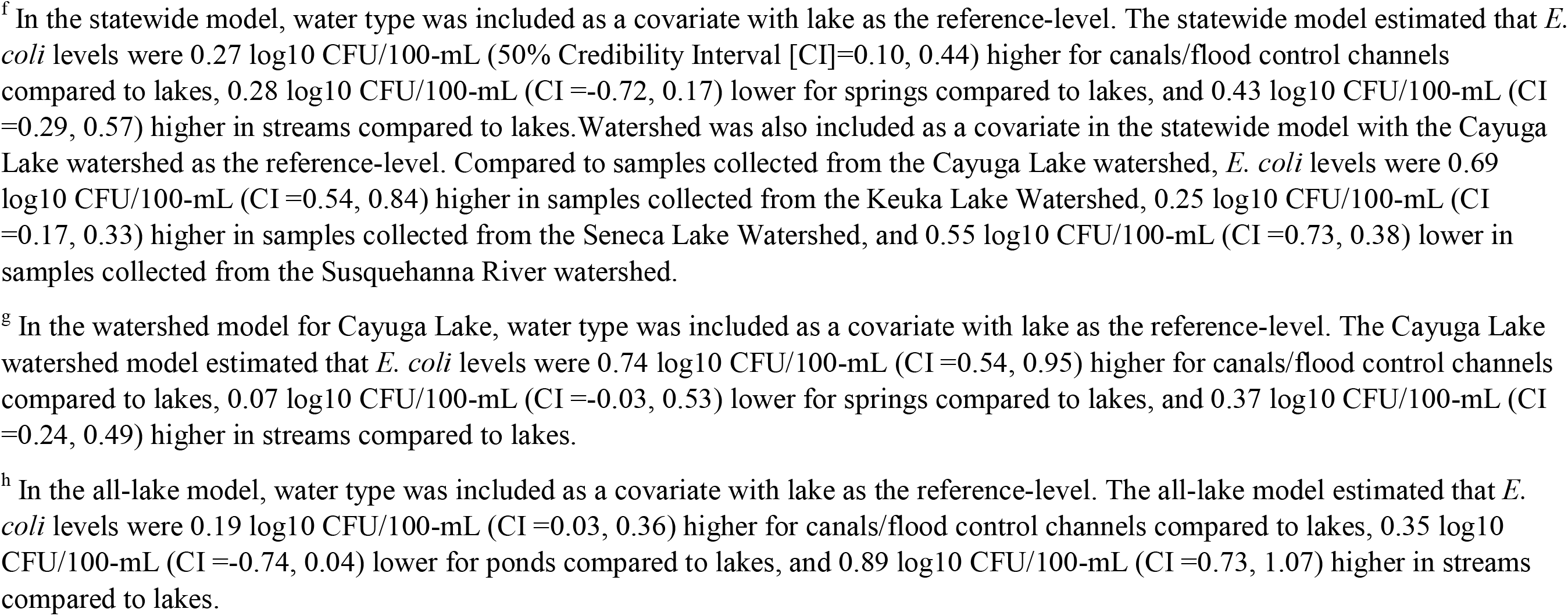
Association between *E. coli* levels and environmental factors according to the multivariable Bayesian models of coregionalization.

**Table S5:**
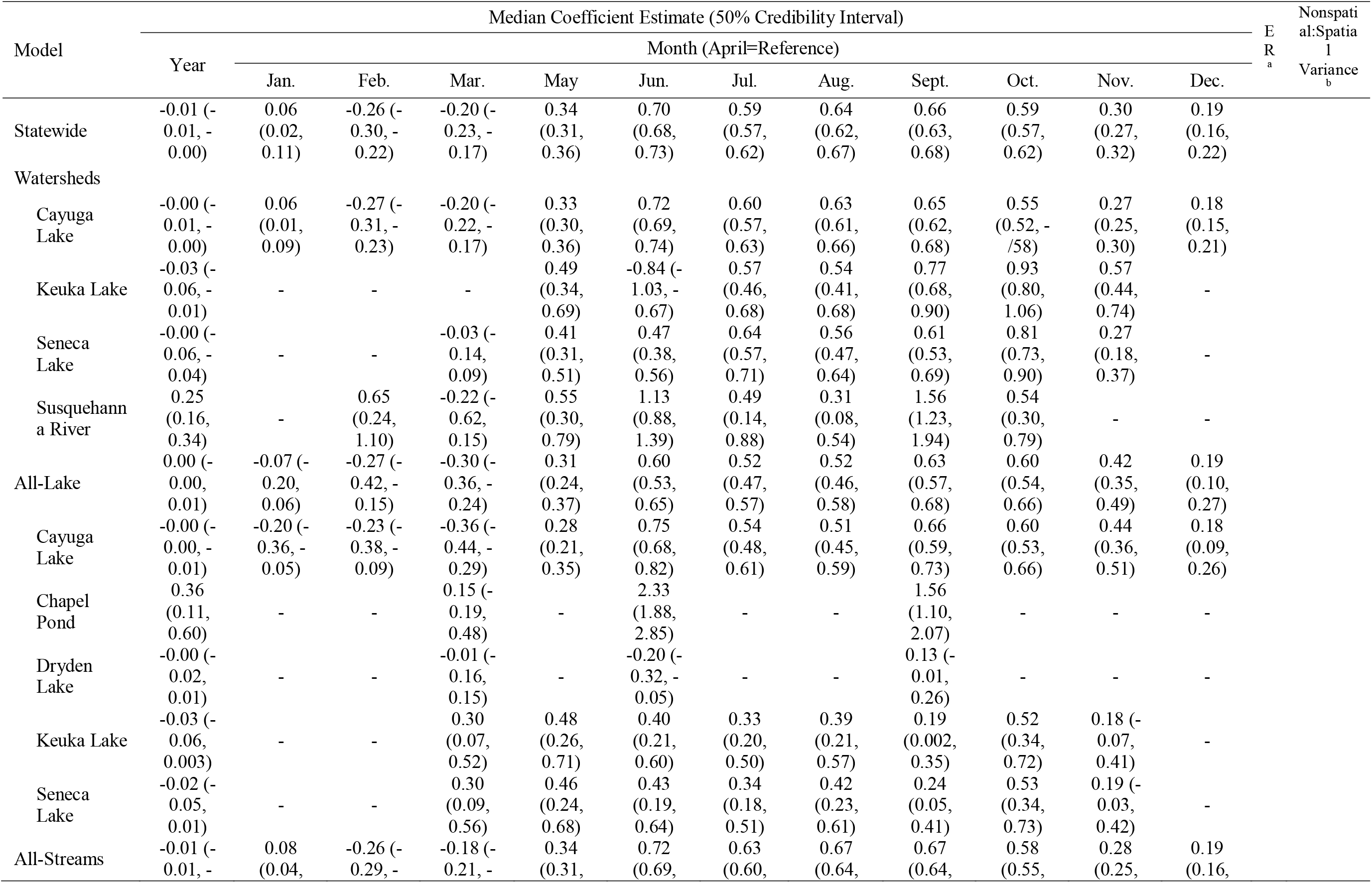
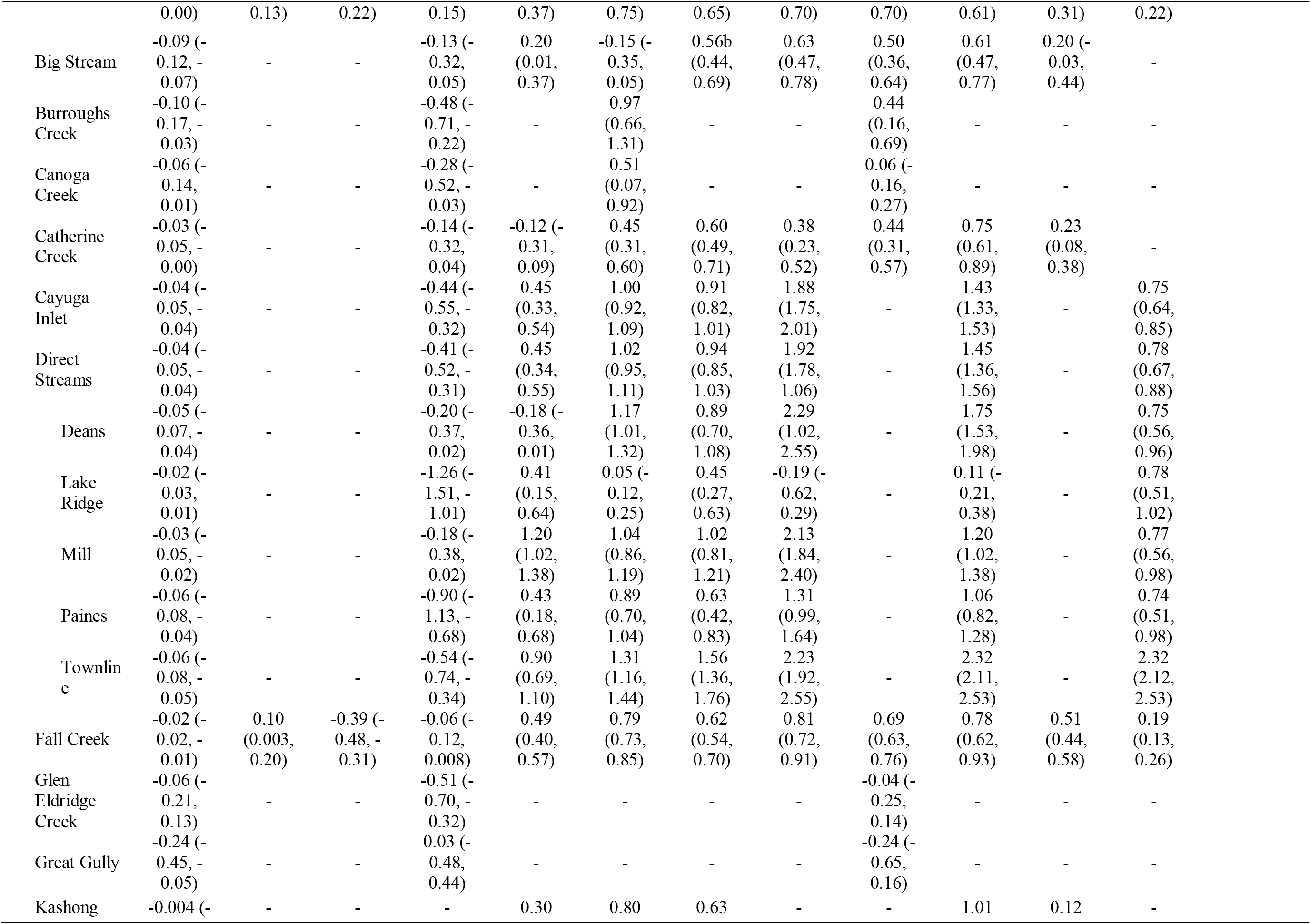
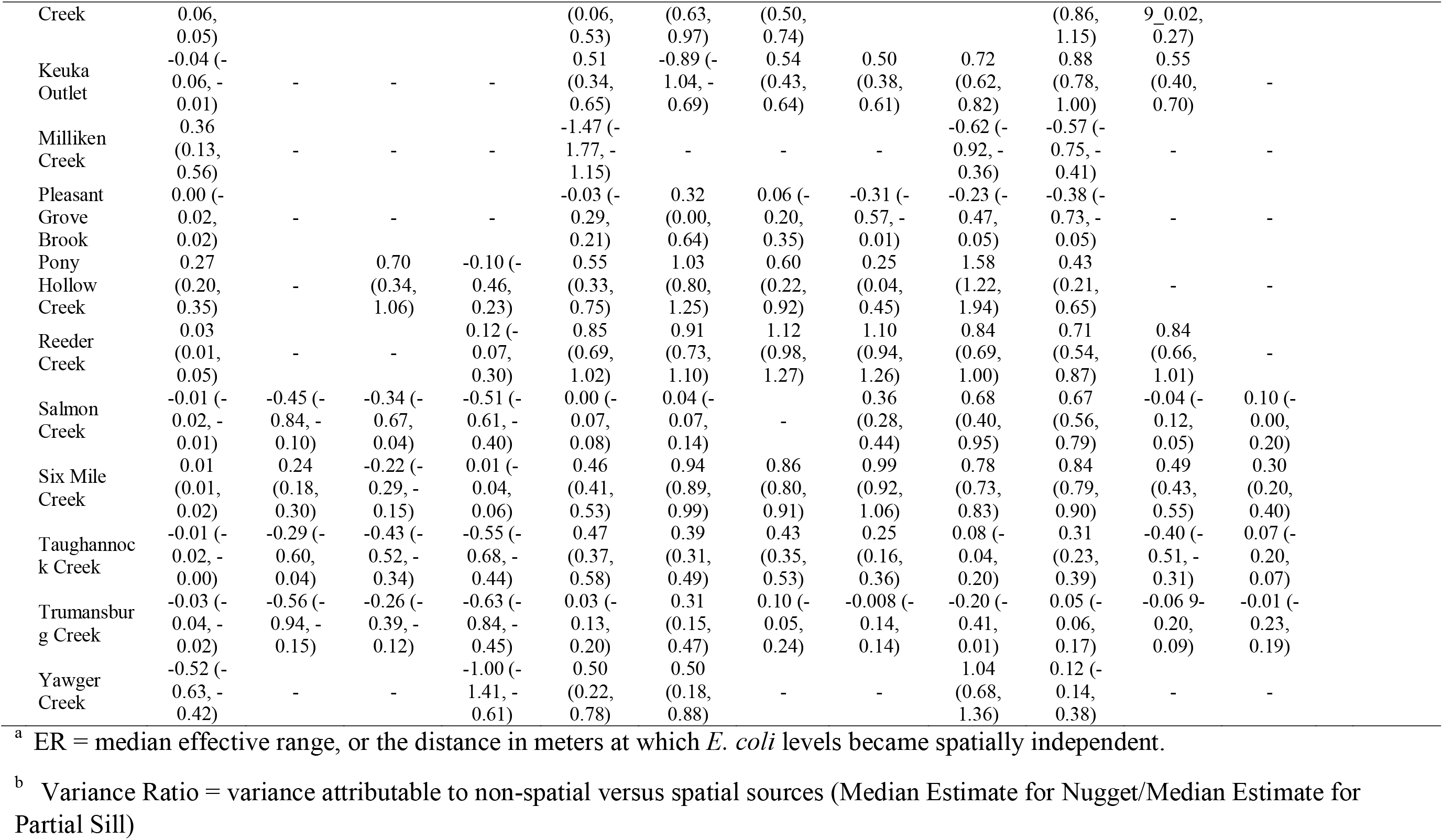
Association between *E. coli* levels and temporal factors according to the multivariable Bayesian models of coregionalization.

**Figure S1:**
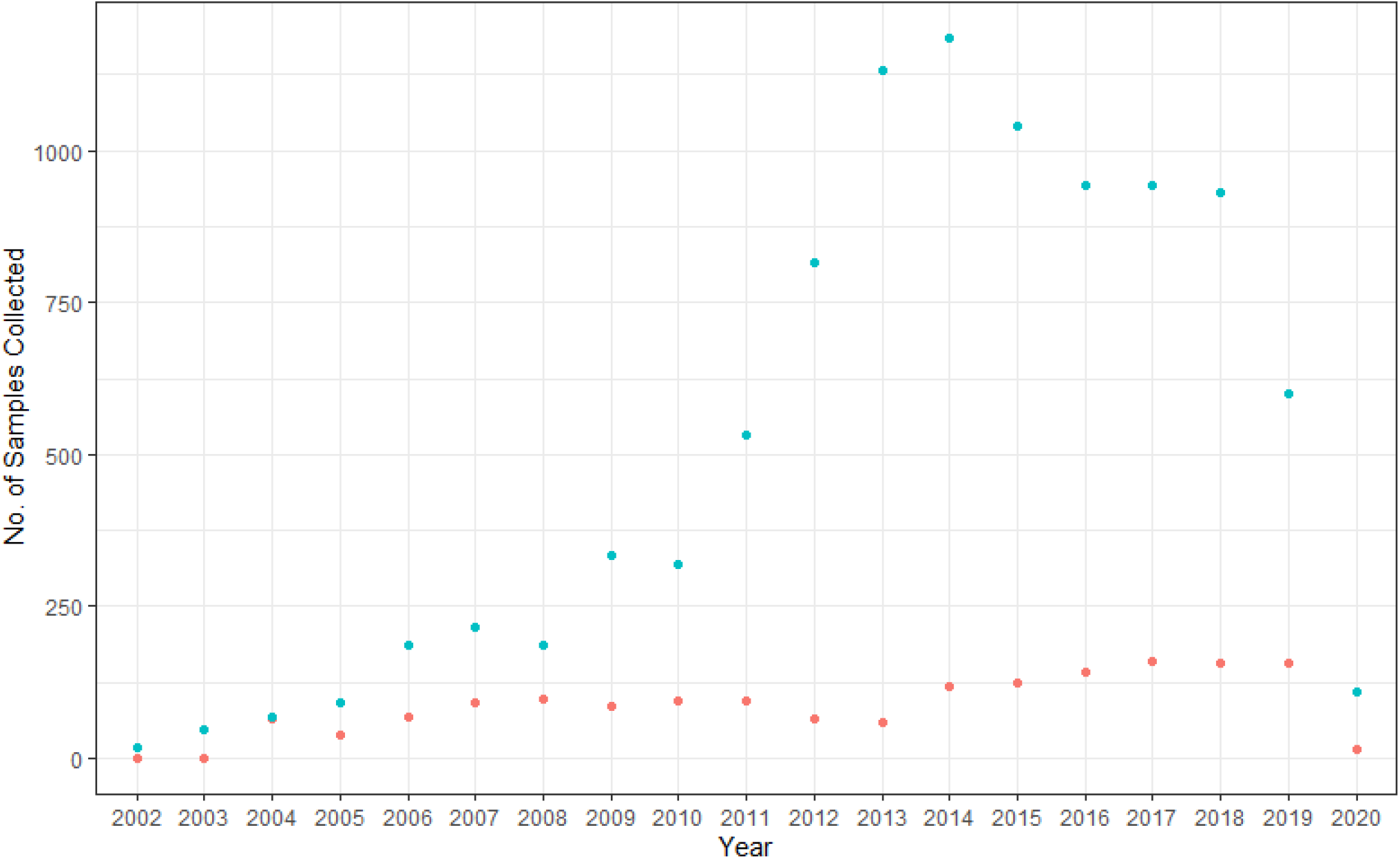
Number of samples collected from lakes and streams each year since collection started. Fewer samples were collected in 2020 than 2019 since data were downloaded in April 2020; the most recent sample was collected on April 6^th^, 2020.

**Figure S2:**
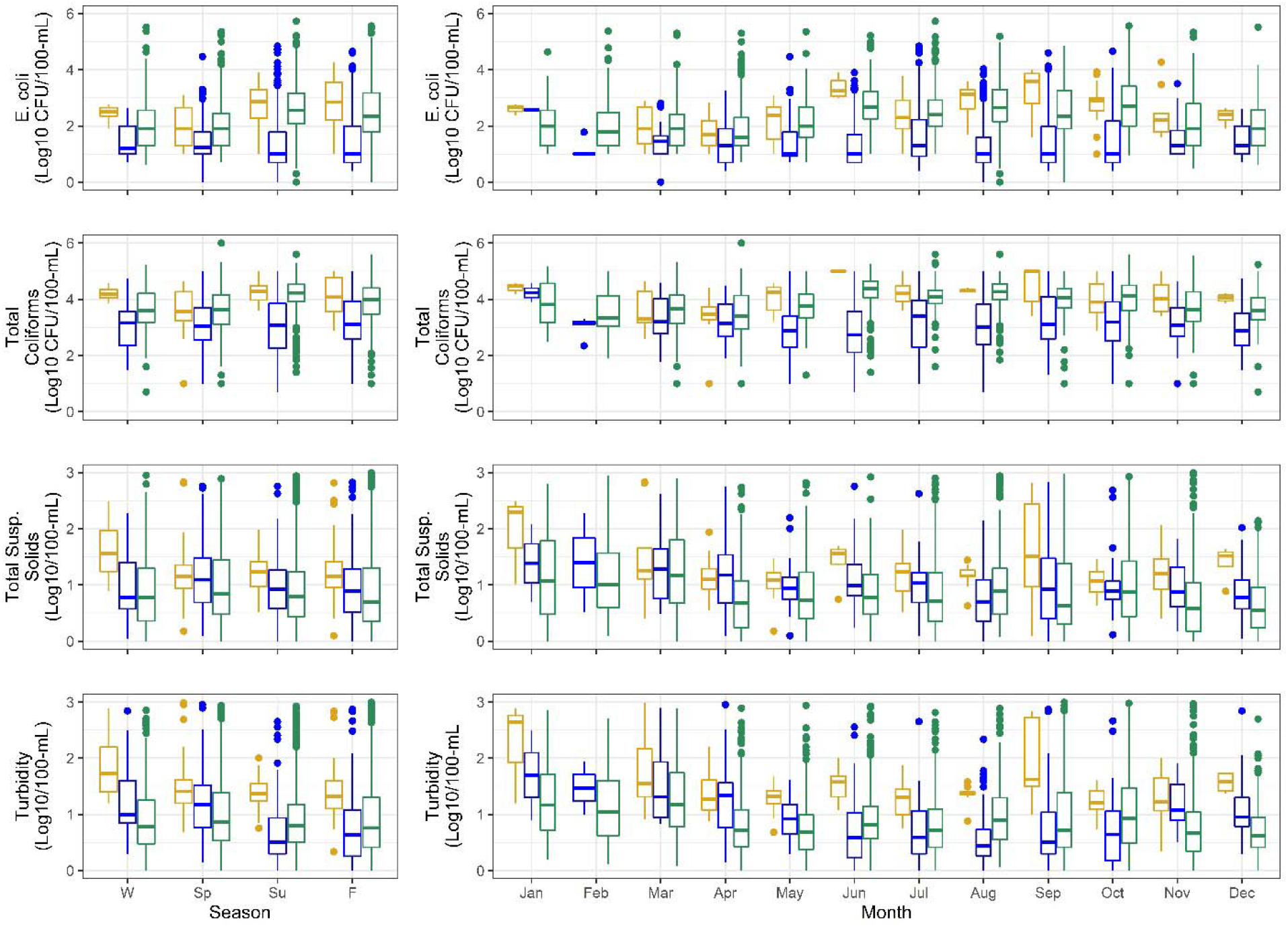
Seasonal (W=winter, Sp=spring, Su=summer, F=fall) and monthly differences in the distribution of log10 *E. coli*, total coliform, total suspended solids, and turbidity in canal, lake, and streams samples.

**Figure S3:**
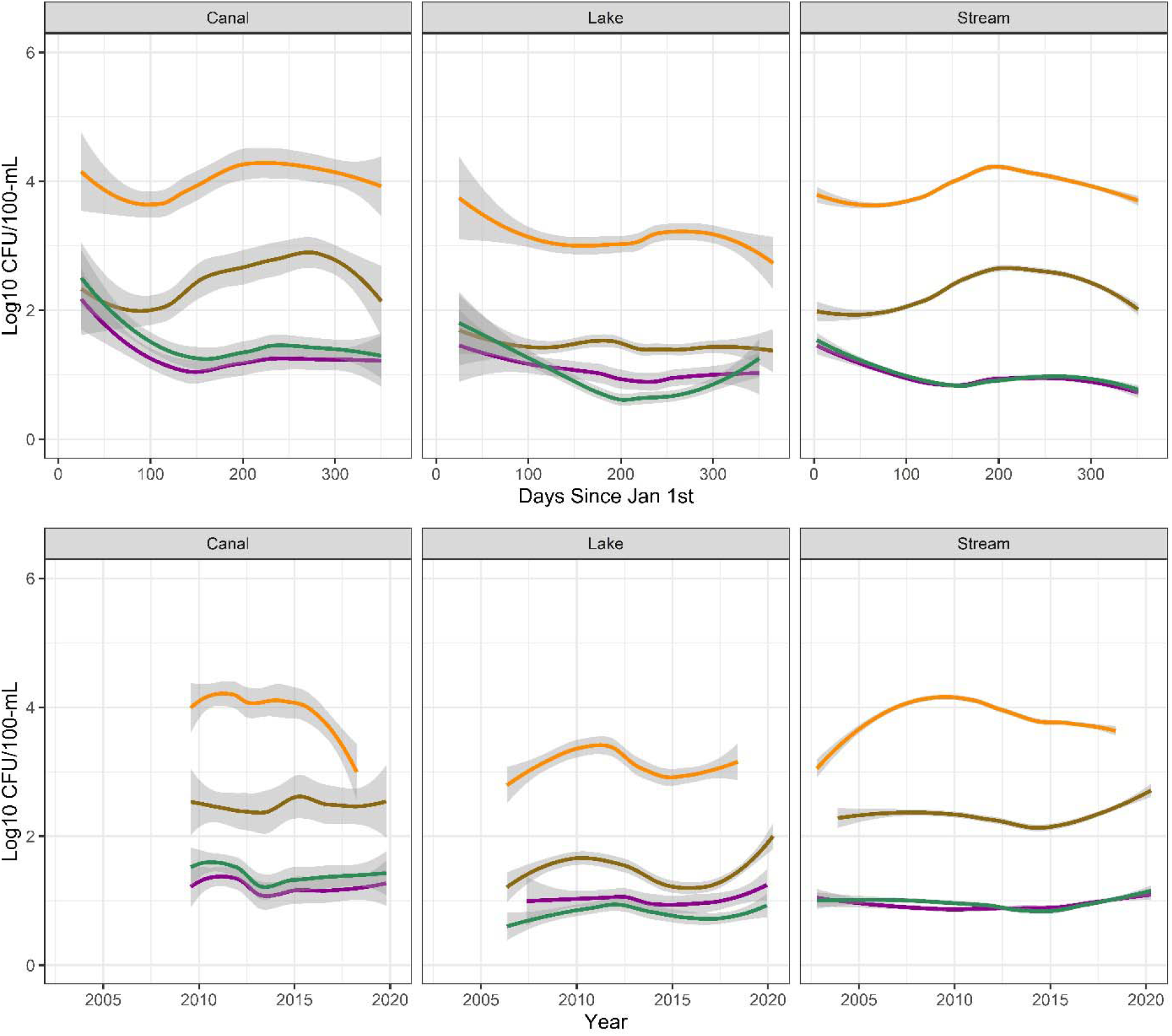
Loess-smoothed lines showing trends in the log10 concentration of ***E. coli***, **total coliforms**, **total suspended solids**, and **turbidity** against days since Jan 1^st^ (i.e., over one year), and over the 18 years samples were collected.

**Figure S4:**
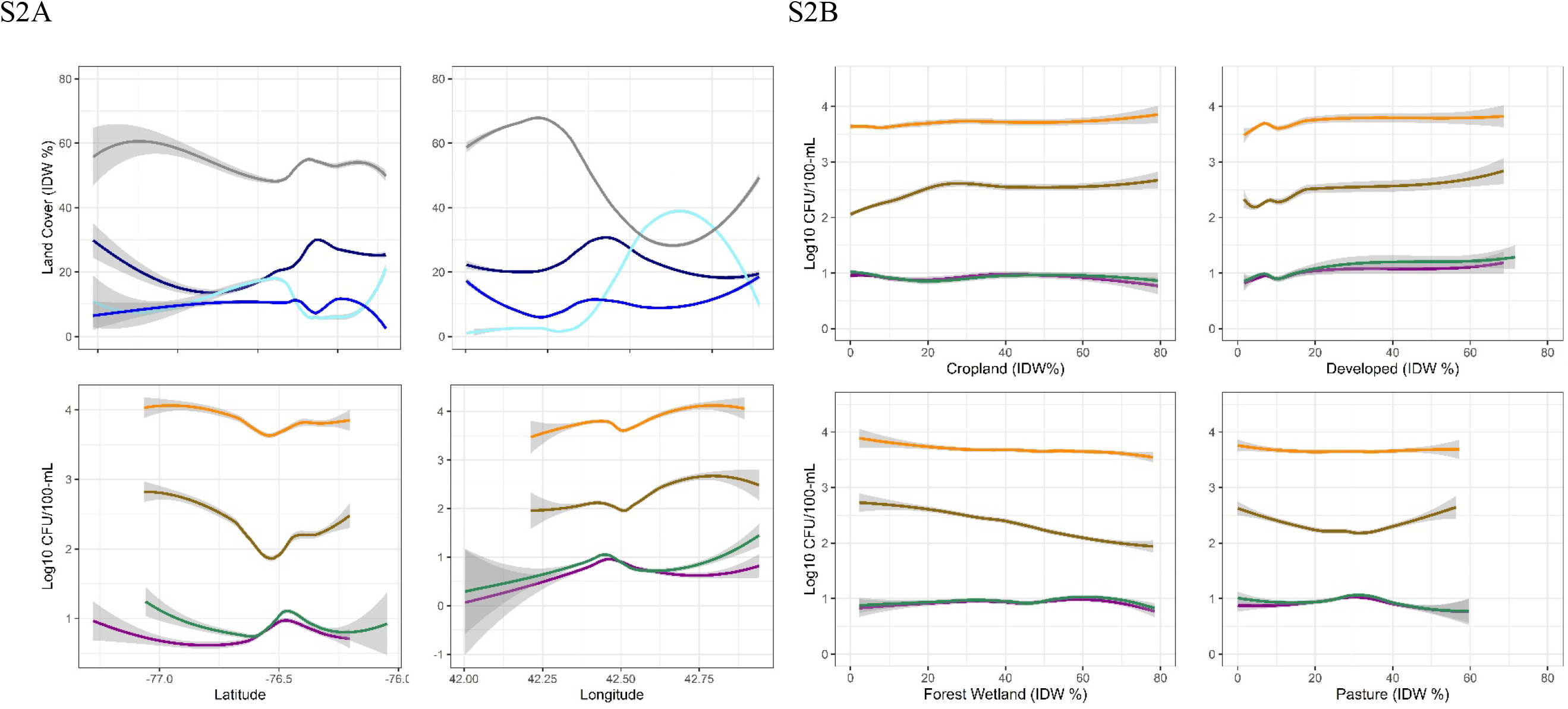
**(S2A)** Loess-smoothed lines showing East-West and North-South trends in (Top Row) the IDW percent of the watersheds under **crop**, **developed**, **forest-wetland**, and **pasture** cover, and (Bottom Row) the log10 concentration of ***E. coli***, **total coliforms**, **total suspended solids**, and **turbidity**. Note that in FigS2A the trends in land cover approx. mirror the trends in water quality. **(S2B)**Loess-smoothed lines show trends in the log10 concentration of ***E. coli*, total coliforms**, **total suspended solids**, and **turbidity** against IDW percent of upstream watershed under crop, developed, forest-wetland, or pasture cover.

**Figure S5:**
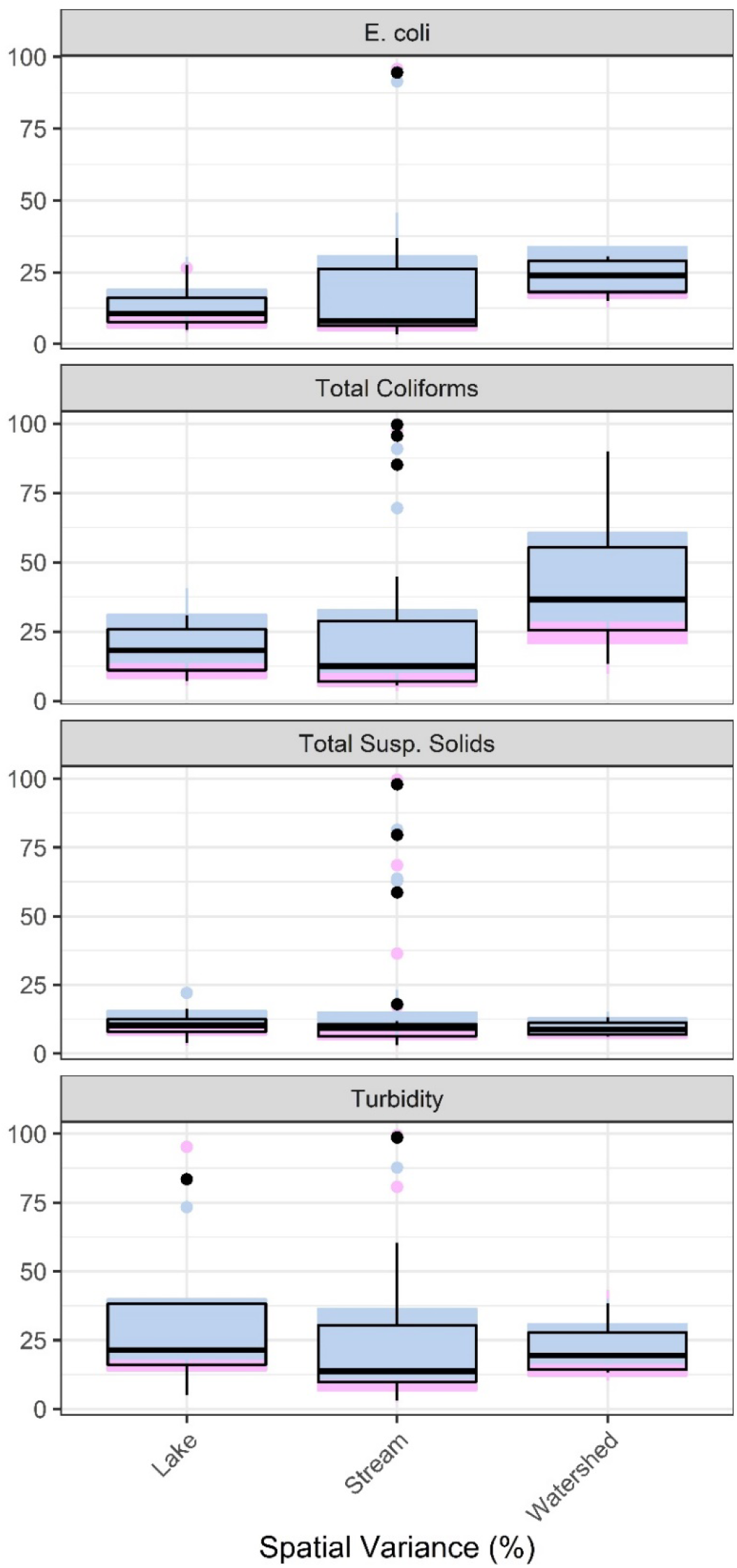
Percent of variance in four water quality parameters that can be attributed to spatial effects in intercept-only Bayesian models of coregionalization for individual lakes (L), streams (S), and HUC-4/Finger Lake watersheds. The percent spatial variance calculated using the 1^st^ and 3^rd^ quartiles, and median values for the partial sill and nugget are shown [% spatial variance (partial sill)/(partial sill+nugget)].

## References

1. Ecological Criteria Division UE. 2009. Review of Published Studies to Characterize Relative Risks from Different Sources of Fecal Contamination in Recreational Water, EPA 822-R-09-001, February 2009.

2. Barton Behravesh C, Mody RK, Jungk J, Gaul L, Redd JT, Chen S, Cosgrove S, Hedican E, Sweat D, Chávez-Hauser L, Snow SL, Hanson H, Nguyen T-A, Sodha S V, Boore AL, Russo E, Mikoleit M, Theobald L, Gerner-Smidt P, Hoekstra RM, Angulo FJ, Swerdlow DL, Tauxe R V, Griffin PM, Williams IT. 2011. 2008 outbreak of Salmonella Saintpaul infections associated with raw produce. N Engl J Med 364:918–927.

3. Ackers M, Mahon B, Leahy E, Goode B, Damrow T, Hayes P, Bibb W, Rice D, Barrett T, Hutwagner L, Griffin P, Slutsker L. 1998. An outbreak of *Escherichia coli* O157:H7 infections associated with leaf lettuce consumption. J Infect Dis 177:1588–1593.

4. Lee S, Levy D, Craun G, Beach M, Calderon R. 2002. Surveillance for Waterborne-Disease Outbreaks --- United States, 1999–2000.

5. Licence K, Oates KR, Synge BA, Reid TMS. 2001. An outbreak of E. coli O157 infection with evidence of spread from animals to man through contamination of a private water supply. Epidemiol Infect 126:135–138.

6. Isaac-Renton JL, Lewis LF, Ong CSL, Nulsen MF. 1994. A second community outbreak of waterborne giardiasis in Canada and serological investigation of patients. Trans R Soc Trop Med Hyg 88:395–399.

7. US FDA. Factors Potentially Contributing to the Contamination of Romaine Lettuce Implicated in the Three Outbreaks of E. coli O157:H7 During the Fall of 2019 | FDA.

8. Food and Drug Administratio. 2018. FDA Investigated Multistate Outbreak of E. coli O157:H7 Infections Linked to Romaine Lettuce from Yuma Growing Region. Center for Food Safety and Applied Nutrition, Washington, D.C.

9. Bottichio L, Keaton A, Thomas D, Fulton T, Tiffany A, Frick A, Mattioli M, Kahler A, Murphy J, Otto M, Tesfai A, Fields A, Kline K, Fiddner J, Higa J, Barnes A, Arroyo F, Salvatierra A, Holland A, Taylor W, Nash J, Morawski BM, Correll S, Hinnenkamp R, Havens J, Patel K, Schroeder MN, Gladney L, Martin H, Whitlock L, Dowell N, Newhart C, Watkins LF, Hill V, Lance S, Harris S, Wise M, Williams I, Basler C, Gieraltowski L. 2019. Shiga Toxin–Producing Escherichia coli Infections Associated With Romaine Lettuce—United States, 2018. Clin Infect Dis.

10. Gorski L, Rivadeneira P, Cooley MB. 2019. New strategies for the enumeration of enteric pathogens in water. Environ Microbiol Rep 1758–2229.12786.

11. Uyttendaele M, Jaykus L-A, Amoah P, Chiodini A, Cunliffe D, Jacxsens L, Holvoet K, Korsten L, Lau M, McClure P, Medema G, Sampers I, Rao Jasti P. 2015. Microbial Hazards in Irrigation Water: Standards, Norms, and Testing to Manage Use of Water in Fresh Produce Primary Production. Compr Rev Food Sci Food Saf 14:336–356.

12. Weller D, Belias A, Green H, Roof S, Wiedmann M. 2020. Landscape, Water Quality, and Weather Factors Associated With an Increased Likelihood of Foodborne Pathogen Contamination of New York Streams Used to Source Water for Produce Production. Front Sustain Food Syst 3:124.

13. Weller D, Brassill N, Rock C, Ivanek R, Mudrak E, Roof S, Ganda E, Wiedmann M. 2020. Complex Interactions Between Weather, and Microbial and Physicochemical Water Quality Impact the Likelihood of Detecting Foodborne Pathogens in Agricultural Water. Front Microbiol 11.

14. Bell RL, Kase JA, Harrison LM, Balan K V., Babu U, Chen Y, Macarisin D, Kwon HJ, Zheng J, Stevens EL, Meng J, Brown EW. 2021. The Persistence of Bacterial Pathogens in Surface Water and Its Impact on Global Food Safety. Pathog 2021, Vol 10, Page 1391 10:1391.

15. ANZECC. 2000. Guidelines for Fresh and Marine Water Quality, 1st ed. Australian and New Zealand Environment and Conservation Council, Agriculture and Resource Management Council of Australia and New Zealand, Auckland, New Zealand, Australia and New Zealand.

16. US FDA. 2015. Standards for the Growing, Harvesting, Packing, and Holding of Produce for Human Consumption, Food Safety Modernization Act. United States.

17. Environmental Protection Agency. 2012. Recreational water quality criteria. Washington, D.C., United States.

18. Health Canada. 2012. Guidelines for Canadian Recreational Water Quality Third Edition, 3rd ed. Ottawa, Canada, Canada.

19. UK EA. Bathing Water Quality.

20. EU Parliament. 2006. Bathing Water Quality Directive. Directive 2006/7/ECOfficial Journal of the European Union.

21. SA DWAF. 1996. Water Quality Guidelines. South Africa.

22. FDA. 2013. United States Food and Drug Administration. Standards for the growing, harvesting, packing, and holding of produce for human consumption.

23. California Leafy Greens Marketing Agreement. 2017. Commodity Specific Food Safety Guidelines for the Production and Harvest of Lettuce and Leafy Greens. California Leafy Green Handler Marketing Board, Sacramento, CA, California, United States.

24. Arizona Leafy Greens Marketing Agreement, California Leafy Greens Marketing Agreement. 2012. Commodity specific food safety guidelines for the production and harvest of lettuce and leafy greens. California Leafy Green Handler Marketing Board, Phoenix, AZ, Arizona, United States.

25. Havelaar AH, Vazquez KM, Topalcengiz Z, Muñoz-Carpena R, Danyluk MD. 2017. Evaluating the U.S. Food Safety Modernization Act Produce Safety Rule Standard for Microbial Quality of Agricultural Water for Growing Produce. J Food Prot 80:1832–1841.

26. McEgan R, Mootian G, Goodridge LD, Schaffner DW, Danyluk MD. 2013. Predicting *Salmonella* populations from biological, chemical, and physical indicators in Florida surface waters. Appl Environ Microbiol 79:4094–4105.

27. Edge TA, El-Shaarawi A, Gannon V, Jokinen C, Kent R, Khan IUH, Koning W, Lapen D, Miller J, Neumann N, Phillips R, Robertson W, Schreier H, Scott A, Shtepani I, Topp E, Wilkes G, van Bochove E. 2012. Investigation of an Environmental Benchmark for Waterborne Pathogens in Agricultural Watersheds in Canada. J Environ Qual 41:21.

28. Truitt LN, Vazquez KM, Pfunter RC, Rideout SL, Havelaar AH, Strawn LK. 2018. Microbial Quality of Agricultural Water Used in Produce Preharvest Production on the Eastern Shore of Virginia. J Food Prot 81:1661–1672.

29. Wall G, Clements D, Fisk C, Stoeckel D, Woods K, Bihn E. 2019. Meeting Report: Key Outcomes from a Collaborative Summit on Agricultural Water Standards for Fresh Produce. Compr Rev Food Sci Food Saf 18:723–737.

30. Francy DS, Stelzer EA, Duris JW, Brady AMGG, Harrison JH, Johnson HE, Ware MW. 2013. Predictive models for Escherichia coli concentrations at inland lake beaches and relationship of model variables to pathogen detection. Appl Environ Microbiol 79:1676–88.

31. Green H, Wilder M, Wiedmann M, Weller D. 2021. Integrative Survey of 68 Non-overlapping Upstate New York Watersheds Reveals Stream Features Associated With Aquatic Fecal Contamination. Front Microbiol 0:2125.

32. Partyka ML, Bond RF, Chase JA, Atwill ER. 2018. Spatiotemporal variability in microbial quality of western US agricultural water supplies: A multistate study. J Environ Qual 47:939–948.

33. Rao G, Eisenberg J, Kleinbaum D, Cevallos W, Trueba G, Levy K, Rao G, Eisenberg JNS, Kleinbaum DG, Cevallos W, Trueba G, Levy K. 2015. Spatial Variability of *Escherichia coli* in Rivers of Northern Coastal Ecuador. Water 7:818–832.

34. Rafi K, Wagner KL, Gentry T, Karthikeyan R, Dube A. 2018. *Escherichia coli* Concentration as a Function of Stream Order and Watershed Size. J Environ Qual 47:949–957.

35. Wyer MD, Kay D, Morgan H, Naylor S, Clark S, Watkins J, Davies CM, Francis C, Osborn H, Bennett S. 2018. Within-day variability in microbial concentrations at a UK designated bathing water: Implications for regulatory monitoring and the application of predictive modelling based on historical compliance data. Water Res X 1:100006.

36. Crosby SC, Spiller NC, Tietz KE, Cooper JR, Fraboni PJ. 2019. Temporal and spatial variability of instream indicator bacteria (Escherichia coli) and implications for water quality monitoring. Environ Monit Assess 191.

37. Buck O, Niyogi DK, Townsend CR. 2004. Scale-dependence of land use effects on water quality of streams in agricultural catchments. Environ Pollut 130:287–299.

38. Hurley T, Mazumder A. 2013. Spatial scale of land-use impacts on riverine drinking source water quality. Water Resour Res 49:1591–1601.

39. Environmental Protection Agency U, of Water O. 2010. Sampling and Consideration of Variability (Temporal and Spatial) For Monitoring of Recreational Waters.

40. Quilliam RS, Clements K, Duce C, Cottrill SB, Malham SK, Jones DL. 2011. Spatial variation of waterborne Escherichia coli – implications for routine water quality monitoring. J Water Health 9:734–737.

41. Weller DL, Love TMT, Belias A, Wiedmann M. 2020. Predictive models may complement or provide an alternative to existing strategies for managing enteric pathogen contamination of Northeastern streams used for produce production. Front Sustain Food Syst 4:561517.

42. ESRI. 2014. ArcGIS Desktop: Release 10.2.2. 10.2.2. Environmental Systems Research Institute, Redlands, CA.

43. R Core Development Team, Team. 2017. R: A Language and Environment for Statistical Computing. 10.1. R Foundation for Statistical Computing, Vienna, Austria.

44. Wei T. 2013. corrplot: Visualization of a correlation matrix. R package version 0.73. 0.73.

45. Bates D, Maechler M, Bolker B, Walker S. 2014. lme4: Linear mixed-effects models using Eigen and S4. R package version 1.1-7. J Stat Softw.

46. Barton K. 2016. MuMIn: Multi-Model Inference. R package version 1.15.6.

47. Harrand S, Weller D, Mercedes Illas-Ortiz P, Wiedmann M, Strawn L. Tracking and modeling of Listeria monocytogenes contamination in spinach fields from planting to harvest.

48. Wickham H. 2009. ggplot2: elegant graphics for data analysis. Springer New York, Dordrecht; New York.

49. Ishii S, Sadowsky MJ. 2008. *Escherichia coli* in the Environment: Implications for Water Quality and Human Health. Microbes Environ 23:101–108.

50. Finley AO, Banerjee S, Carlin BP. 2007. spBayes: An R Package for Univariate and Multivariate Hierarchical Point-referenced Spatial Models. J Stat Softw 19:1–24.

51. Finley AO, Banerjee S, Gelfand AE. 2015. spBayes for Large Univariate and Multivariate Point-Referenced Spatio-Temporal Data Models. J Stat Softw 63:1–28.

52. Plummer M, Best N, Cowles K, Vines K. 2006. CODA: convergence diagnosis and output analysis for MCMC - Open Research Online. R News 6.

53. Partyka ML, Bond RF, Chase JA, Atwill ER. 2018. Spatial and temporal variability of bacterial indicators and pathogens in six California reservoirs during extreme drought. Water Res 129:436–446.

54. Dorner SM, Anderson WB, Gaulin T, Candon HL, Slawson RM, Payment P, Huck PM. 2007. Pathogen and indicator variability in a heavily impacted watershed. J Water Health 5:241–257.

55. Pachepsky YA, Allende A, Boithias L, Cho K, Jamieson R, Hofstra N, Molina M. 2018. Microbial Water Quality: Monitoring and Modeling. J Environ Qual 47:931–938.

56. R.D. Harmel, R. Karthikeyan, T. Gentry, R. Srinivasan. 2010. Effects of Agricultural Management, Land Use, and Watershed Scale on E. coli Concentrations in Runoff and Streamflow. Trans ASABE 53:1833–1841.

57. Wiens JA. 1989. Spatial scaling in ecology. Funct Ecol 3:385–197.

58. Reicherts JD, Emerson CW. 2009. Monitoring bathing beach water quality using composite sampling. Environ Monit Assess 2009 1681 168:33–43.

59. Kinzelman JL, Dufour AP, Wymer LJ, Rees G, Pond KR, Bagley RC. 2009. Comparison of Multiple Point and Composite Sampling for Monitoring Bathing Water Quality. http://dx.doi.org/101080/07438140609353887 22:95–102.

60. Lothrop N, Bright KR, Sexton J, Pearce-Walker J, Reynolds KA, Verhougstraete MP. 2018. Optimal strategies for monitoring irrigation water quality. Agric Water Manag 199:86–92.

61. Reicherts J. 2008. Using Composite Sampling Techniques to Monitor Bathing Beach Water Quality in Kalamazoo County, Michigan. Masters Theses.

62. Bertke EE. 2007. Composite Analysis for Escherichia coli at Coastal Beaches. J Great Lakes Res 33:335–341.

63. Nagels JW, Davies-Colley RJ, Donnison AM, Muirhead RW. 2002. Faecal contamination over flood events in a pastoral agricultural stream in New Zealand. Water Sci Technol 45:45–52.

64. Muirhead RW, Davies-Colley RJ, Donnison AM, Nagels JW. 2004. Faecal bacteria yields in artificial flood events: quantifying in-stream stores. Water Res 38:1215–1224.

65. Zhou K, Sassi HP, Morrison CM, Duan JG, Gerba CP. 2017. Resuspension of Escherichia coli and MS2 Bacteriophage from Bed Sediment in Irrigation Canals. J Irrig Drain Eng 143.

66. Fluke J, González-Pinzón R, Thomson B. 2019. Riverbed Sediments Control the Spatiotemporal Variability of E. coli in a Highly Managed, Arid River. Front Water 1:4.

67. Crump BC, Peterson BJ, Raymond PA, Amon RMW, Rinehart A, McClelland JW, Holmes RM. 2009. Circumpolar synchrony in big river bacterioplankton. Proc Natl Acad Sci U S A 106:21208–21212.

68. Whitlock JE, Jones DT, Harwood VJ. 2002. Identification of the sources of fecal coliforms in an urban watershed using antibiotic resistance analysis. Water Res 36:4273–4282.

69. Varness KJ, Pacha RE, Lapen RF. 1978. Effects of dispersed recreational activities on the microbiological quality of forest surface water. Appl Environ Microbiol 36:95–104.

70. Christensen H, Pacha R, Varness K, Lapen R. 1978. Human use in a dispersed recreation area and its effect on water quality, p. 107–119. *In* Proceedings, recreational impact on wildlands. U.S. Department of Agriculture, Seattle, WA.

71. Dasher DH, Urban L V, Dvoracek MJ, Fish EB, Abstract A. 1981. Effects of Recreation on Water Quality in Guadalupe Mountains National Park.

72. Hendry GS, Leggatt EA. 1982. Some effects of shoreline cottage development on lake bacteriological water quality. Water Res 16:1217–1222.

73. Flack JE, Medine AJ, Hansen-Bristow KJ. 1988. Stream Water Quality in a Mountain Recreation Area. Mt Res Dev 8:11.

74. Sliva L, Williams DD. 2001. Buffer zone versus whole catchment approaches to studying land use impact on river water quality. Water Res 35:3462–3472.

75. Staley ZR, Grabuski J, Sverko E, Edge TA. 2016. Comparison of microbial and chemical source tracking markers to identify fecal contamination sources in the Humber River (Toronto, Ontario, Canada) and associated storm water outfalls. Appl Environ Microbiol 82:6357–6366.

76. Sauer EP, VandeWalle JL, Bootsma MJ, McLellan SL. 2011. Detection of the human specific Bacteroides genetic marker provides evidence of widespread sewage contamination of stormwater in the urban environment. Water Res 45:4081–4091.

77. Garcia-Armisen T, Servais P. 2007. Respective contributions of point and non-point sources of E. coli and enterococci in a large urbanized watershed (the Seine river, France). J Environ Manage 82:512–518.

